# Proteomics profiling of human synovial fluid suggests global increased protein interplay in early-osteoarthritis (OA) and lost in late-stage OA

**DOI:** 10.1101/2020.09.04.279422

**Authors:** Neserin Ali, Aleksandra Turkiewicz, Velocity Hughes, Elin Folkesson, Jon Tjörnstand, Paul Neuman, Patrik Önnerfjord, Martin Englund

## Abstract

The underlying molecular mechanisms in osteoarthritis (OA) development are largely unknown. This study explores the proteome and the pairwise interplay of proteins on a global level in synovial fluid from patients with late-stage knee OA (arthroplasty), early knee OA (arthroscopy due to degenerative meniscal tear) and from deceased controls without knee OA.

Synovial fluid samples were analyzed using state-of-the-art mass spectrometry with data-independent acquisition. The differential expression of the proteins detected was clustered and evaluated with data mining strategies and a multilevel model. Group-specific slopes of associations were estimated between expressions of each pair of identified proteins to assess the co-expression (i.e. interplay) between the proteins in each group.

More proteins were increased in early-OA vs controls than late-stage OA vs controls. For most of these proteins, the fold changes between late-stage OA vs controls and early stage OA vs controls were remarkably similar suggesting potential involvement in the OA process. Further, for the first time this study illustrated distinct patterns in protein co-expression suggesting that the global interplay between the protein machinery is increased in early-OA and lost in late-stage OA. Further efforts should probably focus on earlier stages of the disease than previously considered.

## Introduction

Osteoarthritis (OA), a degenerative joint disease, is one of the most common chronic health conditions and a leading cause of pain and disability among adults ^1,2^. Signs of OA are present in 5 percent of the population between the age of 35-54 years, and the occurrence of the disease increases with age ^3,4^. Despite the high burden of OA, its pathogenesis remains unclear, and no true biological treatment exists on the global market ^2,5^.

The classic diagnosis of OA relies on the presence of clinical symptoms in combination with radiographic findings of joint degeneration, which are seen at a relatively late stage of the disease ^6^. Thus, the search for reliable biomarkers to diagnose OA and to assess its risk of progression is at full speed ^7^. Discovery of OA biomarkers would be useful in identifying individuals with early disease, e.g. for recruitment to clinical trials, as well as for supplementary monitoring of effects of therapies aimed at disease modification. Finding good biomarkers representing the progression of OA goes hand in hand via exploring the pathogenesis. Former studies to disentangle the molecular pathogenesis of OA or to identify biomarkers have typically focused on specific molecules or signalling pathways ^7–9^. However, the “single biomarker strategy” has over and over again shown its lack of sensitivity and specificity for progression of the disease ^9–11^. In contrast, the global crosstalk between molecules could be very informative in order to explore and gain deeper insights into the molecular pathogenesis of OA ^12^. While a number of different omics studies have been carried out ^13–19^, they were limited by the technologies used or availability of biological samples. Therefore, the aim of this study was to explore disease stages of OA using a global discovery approach based on state-of-the-art mass spectrometry (MS) instrumentation and methodology. The disease stages were represented by human synovial fluid from 1) patients with late-stage knee OA, 2) patients with degenerative meniscus tears (indicative of early-stage OA), and 3) controls without OA, to explore the differential expression and the pairwise interplay of proteins measured.

## Experimental procedures

### Materials

The micro bicinchoninic acid (BCA) protein assay kit and Nanosep® 30K Omega Centrifugal Devices were purchased from Pall Life Sciences (Ann Arbor MI, USA) and SOLAμ™ Solid Phase Extraction (SPE) HRP 2mg/1ml 96-well plates were purchased from Thermo Fisher Scientific (Rockford IL, USA). Trypsin (sequencing grade) was purchased from Promega Corporation (Madison, WI, USA). Calcium chloride, formic acid (FA), hydrochloric acid, and ammonium acetate were purchased from Merck (Darmstadt, Germany). Dithiothreitol (DTT) and iodoacetamide were purchased from Sigma-Aldrich (St. Louis, MO, USA). Acetonitrile (ACN) was purchased from Sigma-Aldrich (St. Louis MO, USA).

### Experimental design and statistical rationale

Synovial fluid was sampled from 3 different types of subjects from the MENIX biobank at Skåne University Hospital, Lund: *i*) end-stage medial compartment knee OA patients undergoing total knee arthroplasty (n=11 [3 men and 8 women], age range 55-80 years), the arthroplasty synovial fluid selected for this study will hereafter be called late-stage OA synovial. *ii*) knee arthroscopy patients who had a typical degenerative meniscal tear (n=7 [3 men and 4 women], age range 50-64 years), the arthroscopy synovial fluid selected for this study will hereafter be called early-stage OA synovial. and *iii*) human deceased donors (controls), without evidence of tibiofemoral OA or known clinical knee OA (n=13 [5 men and 8 women] age range 19-79 years). All human deceased donor samples were obtained within 48 h post-mortem, and the specimens were frozen at −80°C within 2 h of extraction. To be eligible as controls, the donor menisci were required to be macroscopically intact. Further, the femoral cartilage (the load bearing region) from the medial compartment of the same donors (also in the biobank) were inspected and required the cartilage to be macroscopically intact. The synovial fluid samples were obtained by transcutaneous aspiration immediately prior to the respective procedures. The donor synovial fluid selected for this study will hereafter be called control synovial. All synovial fluids were centrifuged at 1800 rpm for 10 min and supernatants and pellets were separately frozen and stored at −80°C. The synovial fluid samples’ selection criteria from the biobank were also based on no visual signs of blood contamination. Informed consent was taken before collecting the samples for bio-banking. The sample collection and analysis has been proven by the ethical review committee of Lund University (Durs: 2015/39;2016/865; 2019/3239) and carried out in accordance with relevant guidelines and regulations.

### Sample preparation

The total protein content in the synovial fluid sample was determined using a BCA protein assay kit. From each sample 50 μL synovial fluid were mixed with 10 μL MS-safe proteinase inhibitor cocktail and 10 μL hyaluronidase 10 μL / μg protein and further incubated for 3 h at 37°C. Four out of the 7 samples in the early OA samples were diluted with saline solution during sampling of synovial fluid; these samples were adjusted to the mean total protein content calculated from the remaining 3 samples in that group. All samples were depleted of the 7 most abundant proteins with the multiple affinity removal system (MARS Hu7 spin cartridge) according to the manufacturers protocol (Agilent Technologies). Samples were further reduced using 4 mM DTT, with shaking at +56°C for 30 min and alkylated using 16 mM iodoacetamide for 1 h at room temperature in the dark. In order to remove residual salts, the samples were precipitated with 1:9 volume 95% EtOH (with 50mM NaAc) at 4°C overnight. The precipitated samples were centrifuged and the supernatants were removed from the pellets. The samples were dissolved in 0.1 M ammonium bicarbonate and digested with sequencing grade trypsin (Promega) at a protease/protein ratio of 1:50, overnight at 37°C. Following digestion, samples were cleaned up with a 30kDa filter (Pall Life Sciences) and the flow-through was desalted with C18 96 well plates (SOLAμ™ Thermo Fisher Scientific). The samples were further spiked with iRT peptides before analyses with mass spectrometry.

### Instrumentation and data analysis

The samples were analyzed with an EASY-nLC 1000 (Thermo Scientific) coupled to a Thermo Scientific Q-Exactive HFX™ mass spectrometer using data-independent acquisition (DIA). For liquid chromatography, mobile phase A consisted of water containing 0.1% formic acid (FA) and mobile phase B consisted of acetonitrile (ACN) containing 0.1% FA. Peptides were loaded on an Acclaim PepMap® 100 nanoViper pre-column (Thermo Scientific, C18, 3 μm particles, 75 μm i.d. 2 cm long) at 300 nL/min for 125 min using 95% mobile phase A. The peptides were separated on a PepMap® RSLC C18 analytical column (Thermo Scientific, C18, 2 μm particles, 75 μm i.d. 25 cm long) at 300 nL/min using an ACN /formic acid gradient consisting of an initial step of 5–7% B over 5 min followed by 7–20% B over 85 min, 20–30% B over 20 min, 30–90 % B over 5 min, held at 90 % B for 5 min and then equilibrated for 15 min at 3% B, Separation was performed at 45 °C and the total acquisition time was 125 min. DIA settings: method duration 125 min, full scan resolution 120 000, scan range 350-1650 m/z. AGC target 3.0e6, maximum injection time 100 ms, Orbitrap resolution 45 000, AGC 3.0e5 with a variable isolation window 30/ 26/ 22/ 20/ 18/ 20/ 19/ 20/ 21/ 23/ 24/ 26/ 31/ 32/ 37/ 40/ 53/ 66/ 99/ 574 m/z, and normalized collision energy 27 eV. Data-dependent acquisition (DDA) settings: method duration 125 min, mass range 350-1650 m/z., full MS scan resolution 120 000, AGC 3e6, maximum injection time 20 ms, Orbitrap resolution 15 000, AGC target 1.0e5, maximum injection 20ms, normalized collision energy 27 eV. The MS raw data were further analyzed with Spectronaut™Pulsar software (version 12.0.20491.15, Biognosys AG, Switzerland) for protein identification and quantitative data extraction. In total 118 runs were used from both the DDA and DIA files were converted to HTRMS format using HTRMS Converter (Biognosys AG, Switzerland) to generate the spectral library. The human protein fasta files were downloaded from the uniprot database (20190416). Default settings were used with additional modifications: cysteine carbamidomethylation was used as a fixed modification, and deamination, pyro-glutamic acid (N-term Glu to pyroglutamic acid), methionine oxidation, hydroxyproline and acetylation were used as variable modifications. Using Trypsin/P as the specific digestion type with maximum of 2 miss cleavages. A subsequent protein search was conducted in Spectronaut™ Pulsar using the recently created spectral library and the same human database as background proteome. Precursor quantitation was performed at MS2 level, and area under the curve was used as quantitation type.

### Statistical analysis

The quantitative data were extracted using Spectronaut™Pulsar software. The differential expression of proteins between the different groups was analyzed using a linear mixed effects model with the sample group, protein type and their interaction as independent variables and subject as a random effect to account for clustering of proteins within an individual, and transformed differential expression as the outcome ^20^.The protein expression was transformed using logarithm with base 2 before the analysis. Further, the model was adjusted for age, sex and body mass index. Only proteins that had a maximum of one missing value in the early-stage OA group, two missing values in the late-stage OA group and two missing values in the control group were included, to enable enough samples for estimation. Residual diagnostics confirmed adequate model fit. As the sample size of the early-stage OA group was lower than the others due to sample availability, and also because this group had the highest amount of missing data on protein differential expression, a sensitivity analysis where performed where the early stage OA group was, allowing a larger set of proteins to be included in this analysis, 474 proteins (compared to 406 proteins when including all three groups).

### Data visualization and presentation

#### Clustering

Principal component analysis (PCA) was used to cluster the samples based on their protein expression data, after filtering out proteins based on missing values (max one missing value for the early-stage OA group and two for the control and late-stage OA groups, as described above) 406 proteins were included in the analysis. PCA pre-processing included a Pareto scaling using “RFmarkerDetector::paretoscale” functions in R.

#### Protein-protein co-expression

The linear associations between pairs of proteins expression and estimates of the regression slopes for each group were conducted. This by fitting a linear regression model for each pair of proteins (82 418 models in total) with expression of one of the proteins as outcome and expression of the other protein, the group and their interaction as independent variables. The intensities were transformed with logarithm of base two before fitting the models and standardized by subtracting mean protein expression and dividing by the standard deviation, to make the slopes comparable between models.

### Reproducibility and quality control

The samples were run in random order with the insistency of having samples from each group analyzed in the beginning, middle and the end of the sample batch run. A pooled sample was run every 10th sample as a quality control sample. 18 of the samples were run as duplicates to check for reproducibility. The reliability was evaluated by comparing two injections of the same sample and estimating the agreement using the Bland-Altman method ^21,22^. The data was again transformed by the logarithm of base two to stabilize variance.

### Pathway analysis

To evaluate common interactions and pathways of the identified proteins with changed levels, the results were further analyzed with IPA software (Ingenuity Systems, Redwood City, CA, USA, www.ingenuity.com). Differentially expressed proteins were mapped and compared to known pathways, diseases, functions, and connecting regulators using only 406 proteins as a background data set. The data were also evaluated based on upstream regulators with positive z-scores (activating capacity) and negative z-scores (inhibiting capacity). Key regulators are proteins with an earlier established association with the activation or inhibition of many of the identified proteins, but the regulators are not necessarily detected themselves in our MS analysis. Default settings were used except for species, which was set to human. In addition, only experimentally observed relationships were considered. The software was unable to map P69905 protein (Hemoglobin subunit alpha). This protein was therefore not further included in the pathway analysis.

### Validation of mass spectrometry results

Thirty-six out of the 406 proteins that were differentially expressed in our MS data were further validated with the Multiplex® cardiometabolic immunoassay panel (Olink Proteomics, Uppsala, Sweden). The synovial fluid samples were diluted 1:1000 before analysis. Processing, output data quality check, and normalization were performed by Olink Proteomics. All data were delivered as Normalized Protein eXpression (NPX) values on a log_2_ scale. Data values below the level of detection (LOD) were removed from the dataset. Using these data, the main analysis was repeated with linear multilevel models to estimate the differences between the groups in protein expression and relate them to the original results.

## Results

### Label free quantification

In total, 715 proteins were detected and relatively quantified in synovial fluid samples from 31 subjects. After selection of proteins with limited missing values, 474 proteins were eligible for the clustering and statistical analysis for comparing the late-stage OA vs control group and 406 proteins for comparing all 3 groups.

### Clustering

Principal component analysis revealed an unsupervised difference between the individual samples clustering the samples very well in their different groups (Fig 1A.). The protein profile of early-stage OA synovial fluid samples clustered between control and late-stage OA samples. The biplot shows some of the proteins that drive this cluster separation (Fig. 1B). Among these are Chemokine (C-X-C motif) ligand 7 (CXCL7), Decorin (PGS2), S10A8, Zinc fingers and homeoboxes protein 3 (ZHX3).

**Fig 1.**
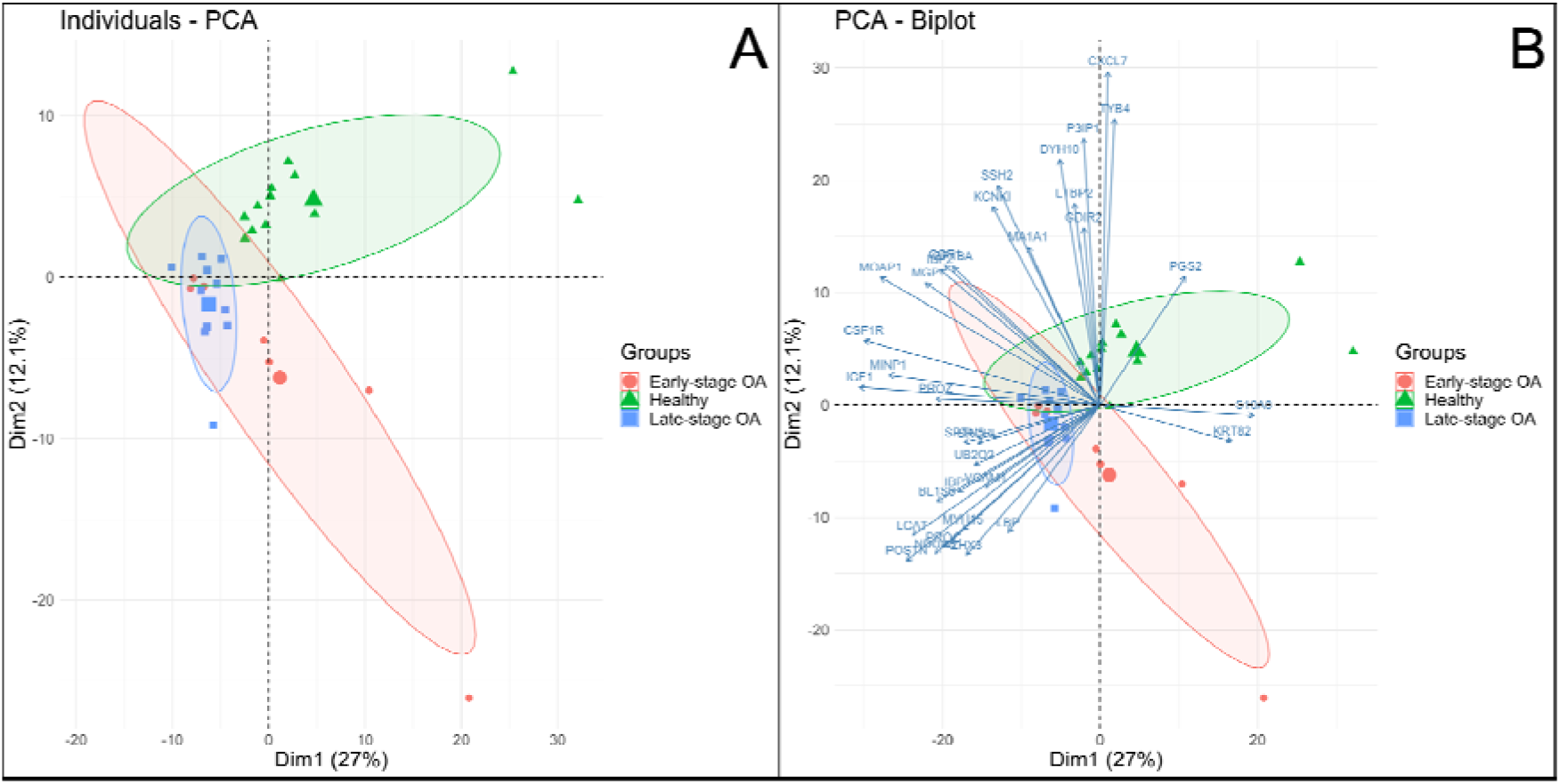
PCA and Biplot; 406 proteins from the label free quantification were used to cluster the different samples. A) Principal component analysis of each sample in each group driven by the label free quantification of the 406 proteins. B) Biplot showing the proteins that drive the separation of the clusters.

### Differentially expressed proteins

Out of 406 proteins analyzed, different patterns of protein expression were identified with over than 200 proteins that where differently expressed between the different groups (Supplemental Fig. S1 and Table S1). Among them 27 proteins (whereof 16 downregulated) with a fold change of 2 or larger in the late-stage OA group compared to control, or in the early-stage OA group compared to control where identified (Fig 2). The most pronounced differences in comparisons with controls were found for PGS2 (fold-change 0.05 with a 95% CI [0.03-0.09] early-stage OA, 0.03 [0.02-0.06] late-stage OA), CXCL7 (0.091 [0.05-0.17] early-stage OA, 0.061 [0.03-0.11] late-stage OA). Glutathione peroxidase 3 (GPX3) (8.85 [4.95-15.8] early-stage OA, 5.91 [3.29-10.63] late-stage OA) and Histidine-rich glycoprotein (HRG) (7.03[3.95-12.51] early-stage OA, 7.79 [4.35-13.95] late-stage OA), where >1 fold changes indicate higher levels in early-stage OA patients than in controls, and vice versa. Inspecting the gradient from the lowest to the highest fold changes in differently expressed proteins (Supplemental Fig. S1), the majority of the proteins that differed between the late-stage OA and control cases were higher in the control samples in comparison to the late-stage OA samples. This observation was even more evident when only comparing the late-stage OA and control samples. When comparing late-stage OA and early-stage OA samples, only 3 proteins were higher in late-stage OA in comparison to early-stage OA – Fibronectin (FINC), 1.99 [1.04-3.83], hemoglobin subunit alpha (HBA), 2.25 [1.17-4.32], Cartilage acidic protein 1 (CRAC1), 1.96 [1.02-3.78], whereas 112 proteins were higher in early-stage OA than in late-stage OA samples. Together, all these results suggest an overall decrease in protein expression in late-stage OA.

**Fig 2.**
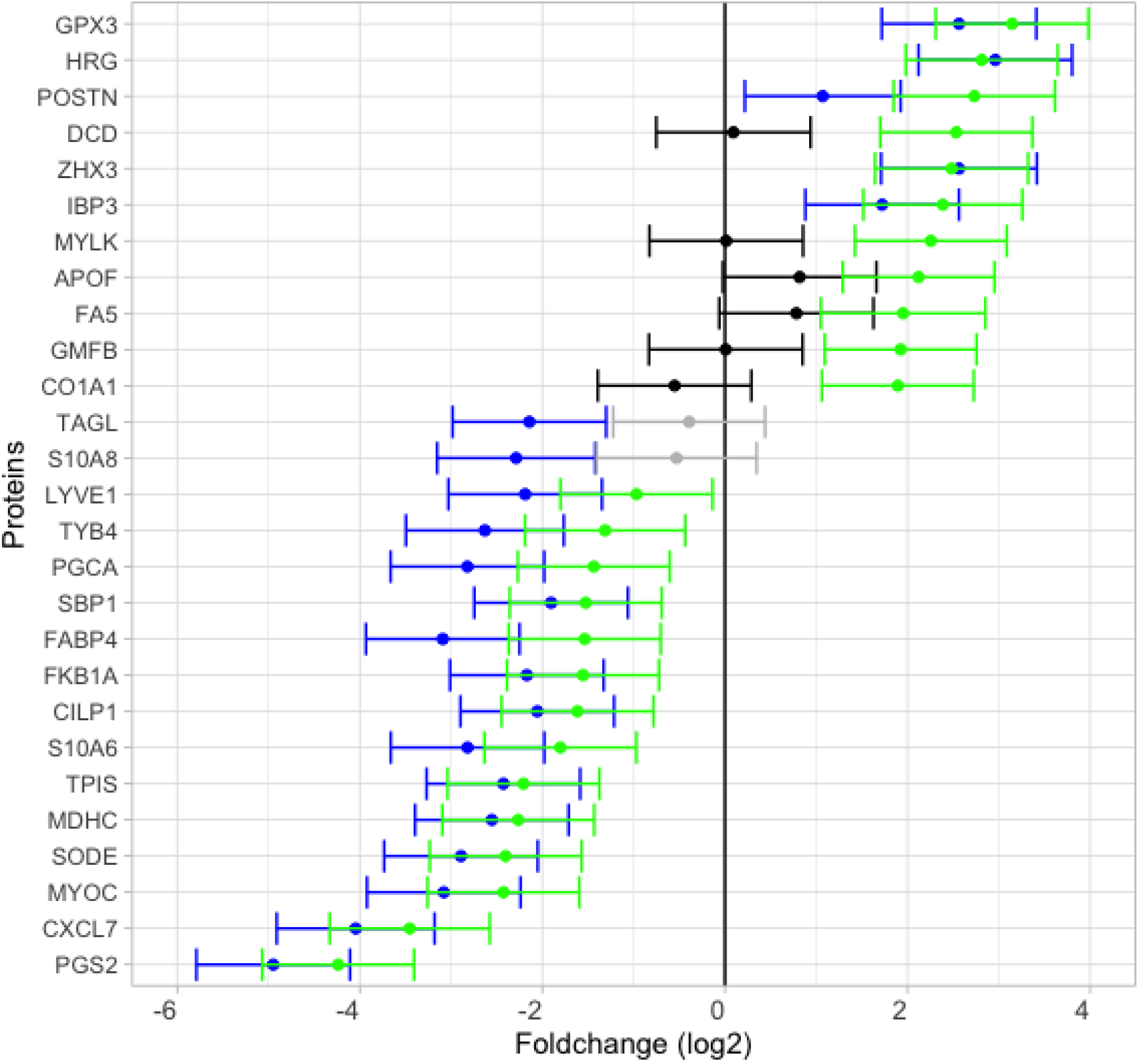
Differentially expressed proteins with an estimated fold change larger than 2. Comparison between early-stage OA vs controls (green and gray bars), Comparison between late-stage OA vs controls (blue and black bars). Blue and green bars represent differentially expressed proteins whose 95% CIs exclude 0.

### Biological relevance

#### Canonical pathways

Pathway analysis revealed that the differentially expressed proteins were connected to multiple canonical pathways (Table 1). The z-scores suggest whether the pathways are activated or inhibited in accordance to the fold changes of the connected proteins.

**Table 1.**
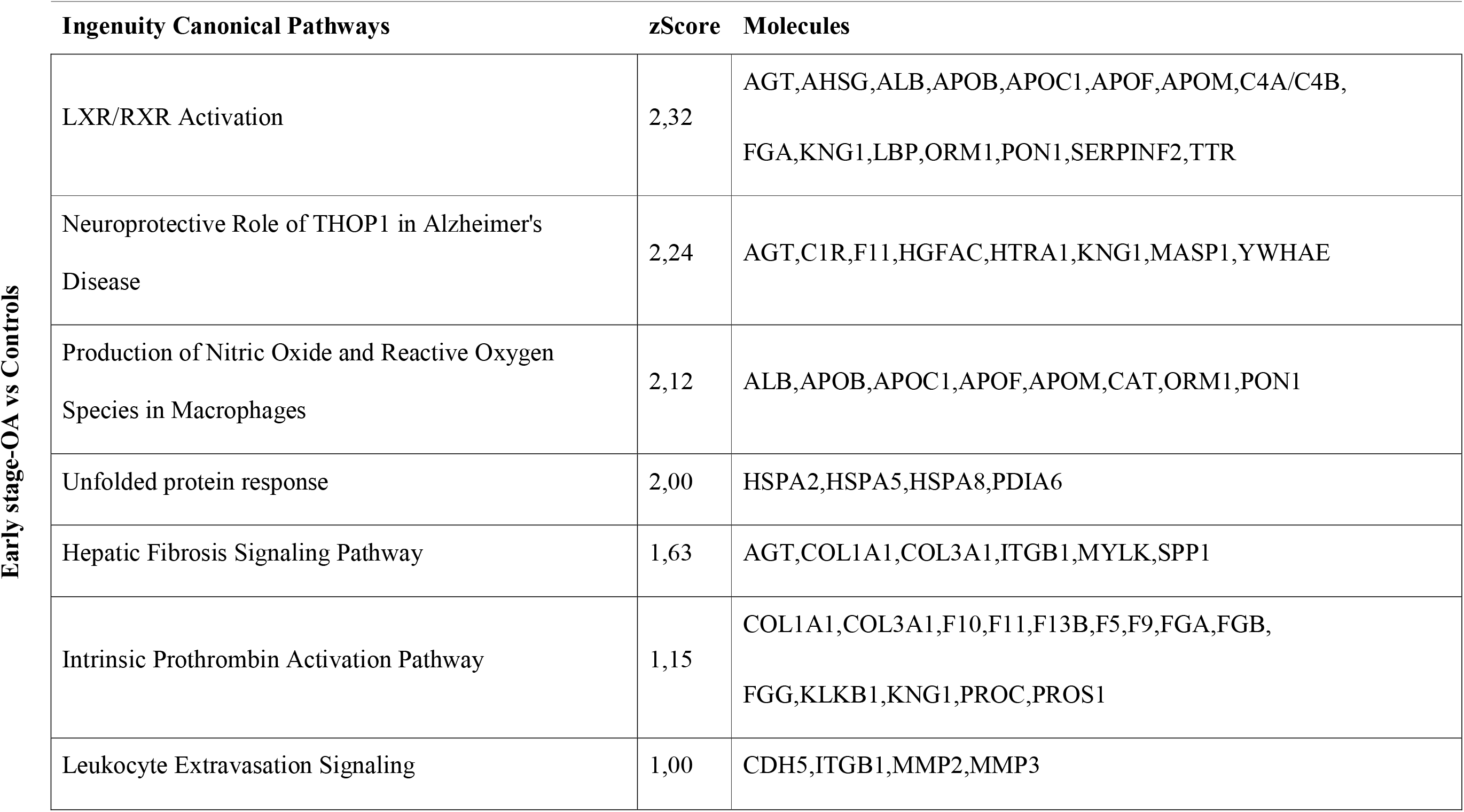
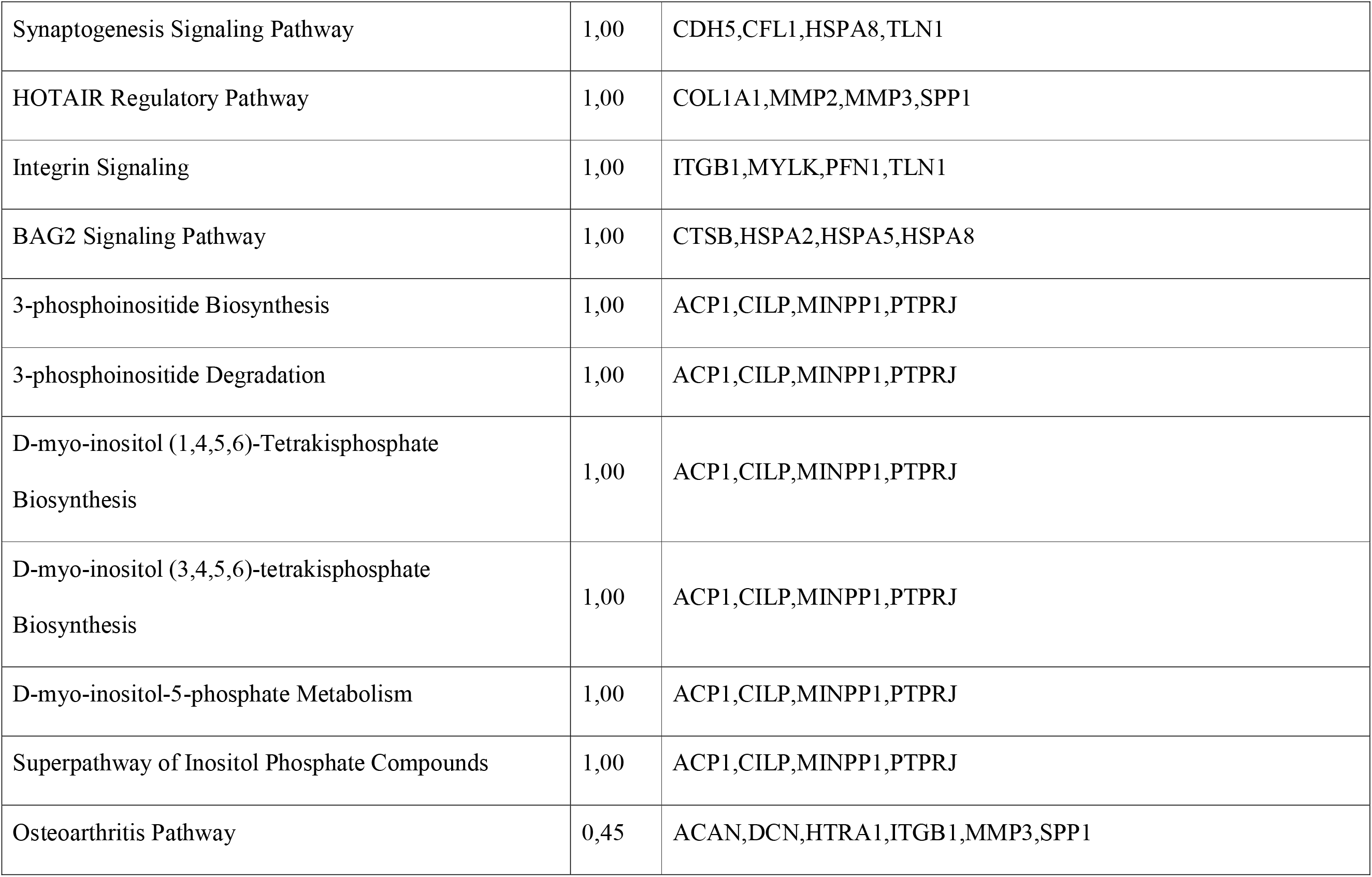
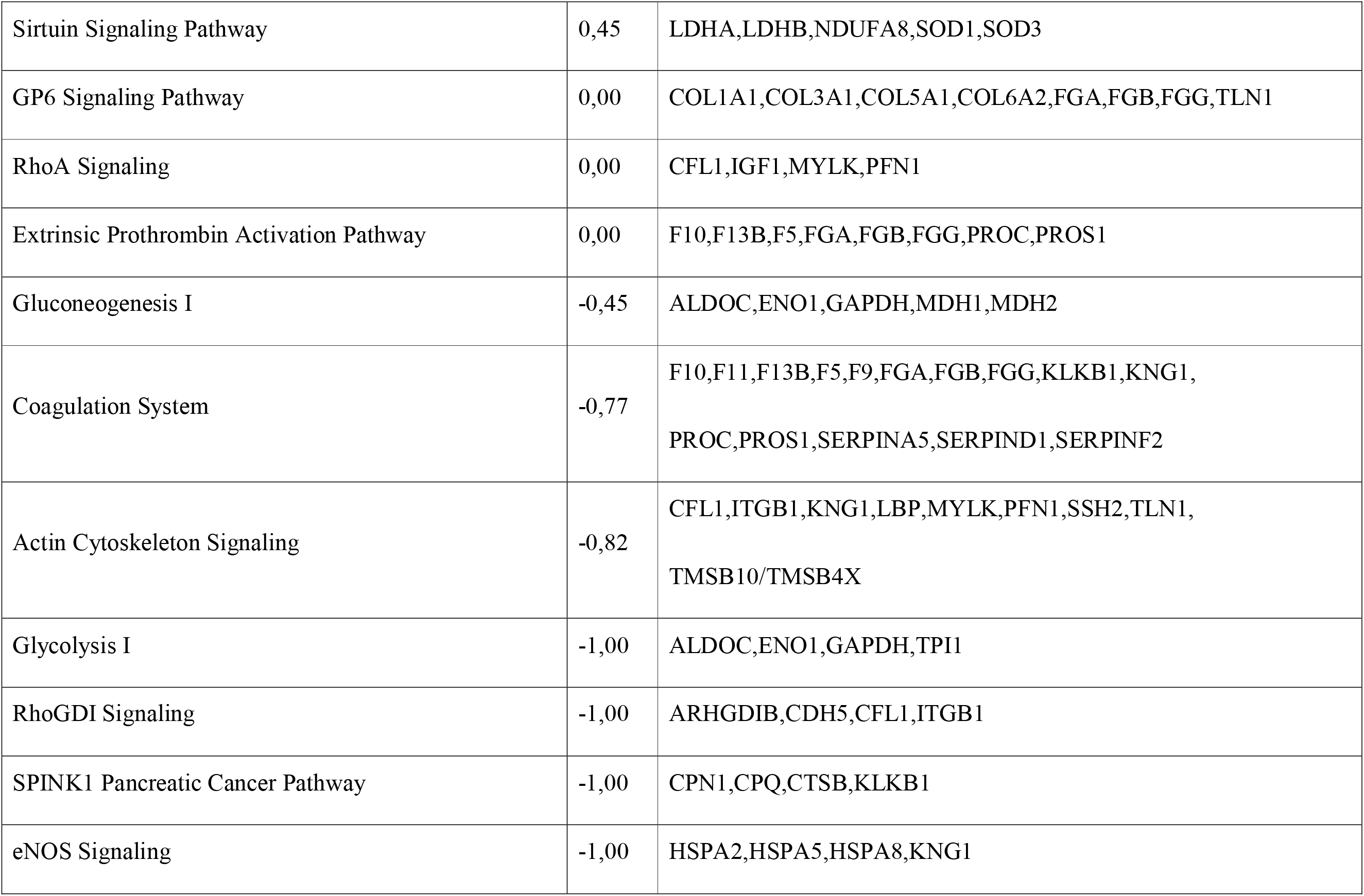
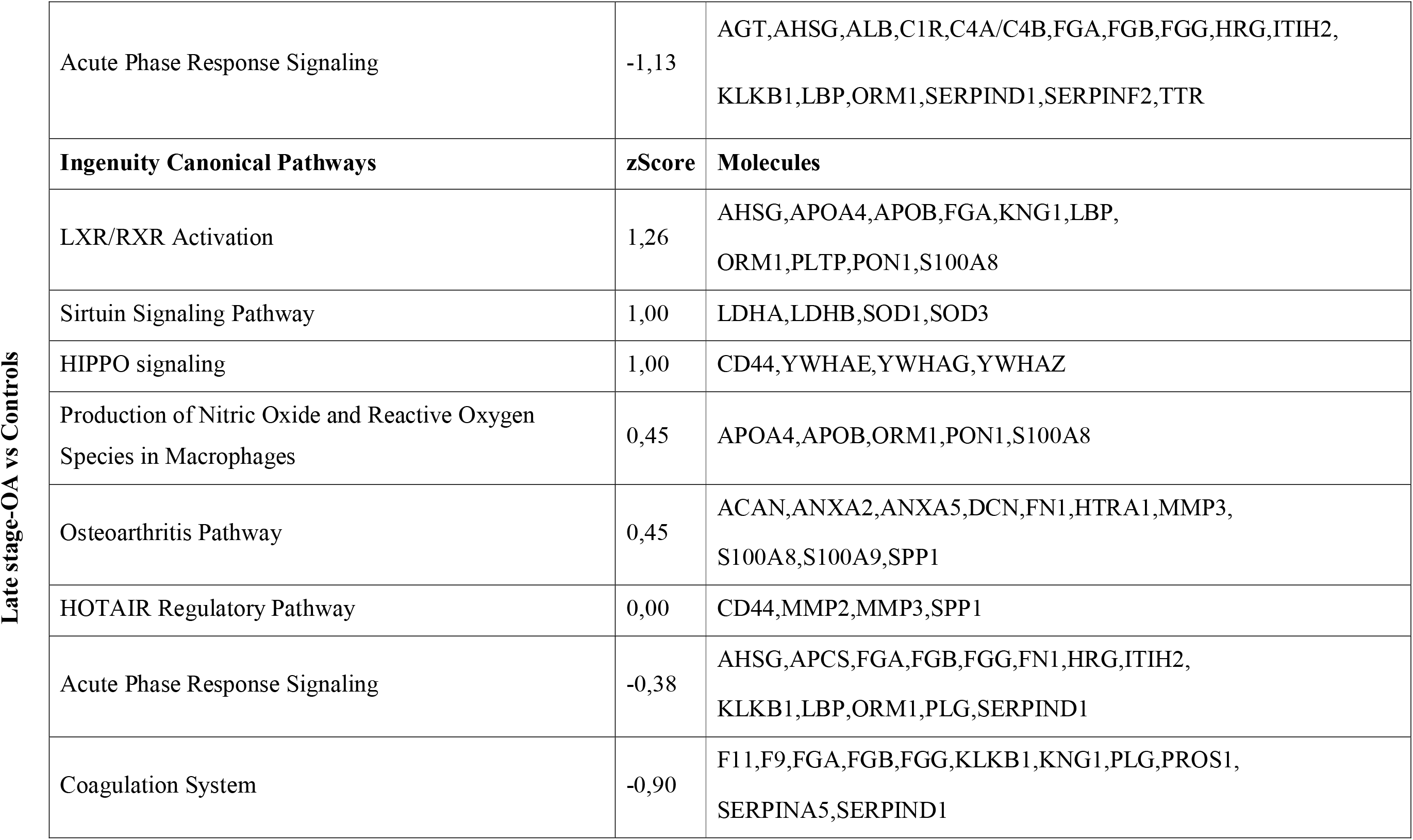
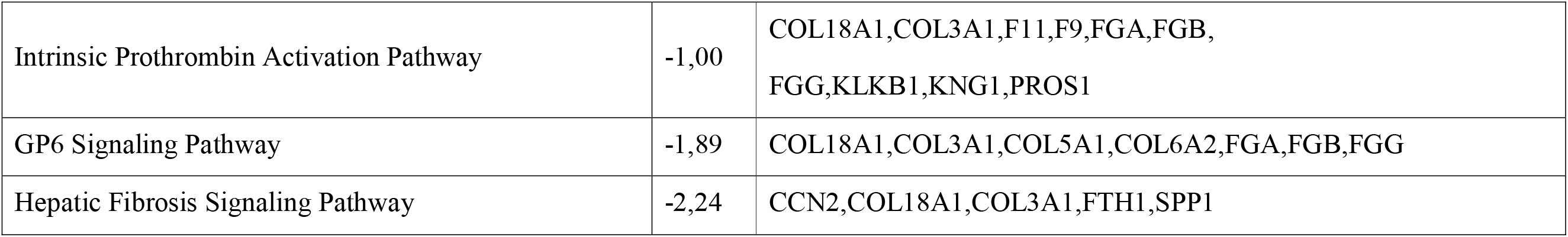
Pathway enrichment analysis. Differentially expressed proteins mapped to canonical pathways. z-Score is the probability score of activation if positive and deactivation if negative.

#### Upstream regulators

Pathway analysis also revealed that the differentially expressed proteins were connected to multiple upstream regulators (Fig. 3). For instance, the transcription factor serum response factor (SRF) was activated in the early-stage OA group in comparison to controls. The analysis also suggested that the transcription factor Y-box-binding protein 1 (YBX1) was activated in the late-stage OA group in comparison to controls.

**Fig 3.**
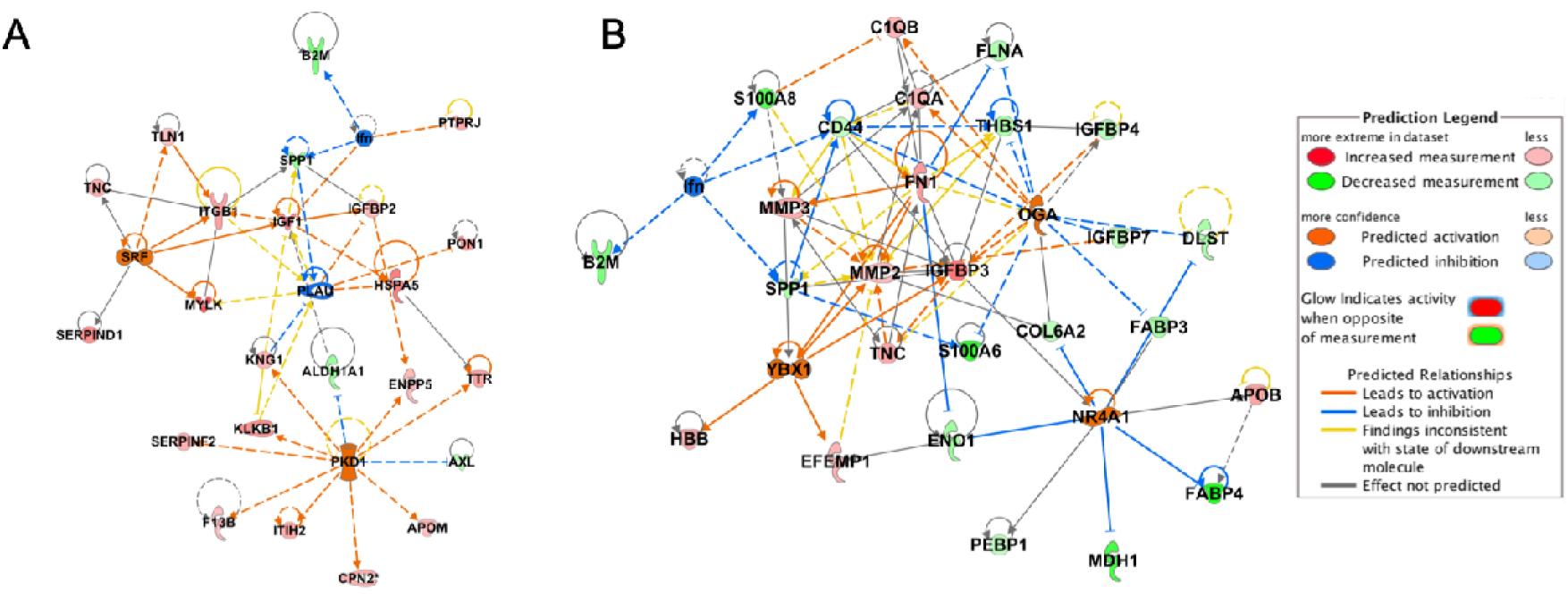
The network of upstream regulators. A) Upstream regulators suggested to be activated or deactivated by the differentially expressed proteins in the early-stage OA group: Ifn, PKD1, PLAU and SRF. B) Upstream regulators suggested to be activated or deactivated from the differentially expressed proteins in the late-stage OA group: Ifn, NR4A1, OGA, YBX1

#### Molecular functions

Pathway analysis revealed that the differentially expressed proteins were connected to multiple diseases and molecular functions (Supplemental Fig. S2 and Table S2). The z-scores suggest whether the corresponding functions are activated or inhibited in accordance to the fold changes of the connected proteins. The overall pattern suggested that several molecular functions such as cellular development and assembly are activated in the early-stage OA group. It also suggested that cellular movement is deactivated in the late-stage OA group.

#### Inflammation

Pathway analysis suggests that the acute phase pathway is inhibited in the early-stage OA and late-stage OA groups in comparison to the controls (Table 1). Leukocyte extravasation signalling is suggested to be activated in the early-stage OA group compared to controls. Additionally, typical markers for inflammation (e.g. CXCL17) were found to be decreased in the early-stage OA and late-stage OA cases in comparison to the controls (Supplemental Table S1).

#### Lipid Metabolism

The pathway analysis suggested a strong activation of the LXR/RXR lipid metabolism pathway in both early-stage OA and late-stage OA groups, with a higher evidence of activation for early-stage OA patients (Table 1).

#### Extracellular matrix (ECM) Organization and damage associated molecular patterns DAMPs

The results revealed multiple ECM proteins that were differentially expressed in the synovial fluid when comparing the different groups to each other (Supplementarl Fig. S1 and Table S1). The results further suggested a cluster of collagens differing between the different groups. CO18A1 (fold-change 0.60 and 95% CI [0.34-1.07] early-stage OA, 0.38 [0.21-0.68] late-stage OA) and CO6A2 (0.47 [0.26-0.85] early-stage OA, 0.53 [0.30-0.96] late-stage OA) had a lower expression in both early-stage OA and late-stage OA cases in comparison to the controls, while CO5A1(3.36 [1.89-5.98] early-stage OA, 2.25 [1.26-4.03] late stage OA) had a higher expression in both early-stage OA and late stage OA cases. CO3A1(0.22 [0.11-0.42]), CO14A1(0.34 [0.17-0.68]) and CO1A1 (0.18 [0.10-0.35]) all had a lower differential expression in the late-stage OA cases compared to early-stage OA cases. Some DAMP proteins did also express differential expression between the different groups, fibrinogen (gamma, beta and alpha) were lower in OA patients in comparison to controls. Tenascin (TENA) (2.12 [1.19-3.77] early-stage OA, 2.13 [1.19-3.82] late-stage OA) was higher in OA patients in comparison to controls.

#### Serine protease activity

Differentially expressed proteins that were higher in OA patients in comparison to controls had serine-type peptidase activity; among them were MMP2 (fold-change 1,91 and 95% CI [1.07-3.40] early-stage OA, 1.79 [1.00-3.20] late-stage OA) and MMP3 (2.38 [1.33-4.24] early-stage OA, 2.08 [1.16-3.72] late-stage OA).

### Associations between pairs of proteins

Protein co-expression data revealed that the control samples have protein pairs that are both positively and negatively co-expressed (mean slope, 0.29), whereas the early-stage OA co-expression revealed an increase in positive co-expression of the same protein pairs (mean slope, 0.48) (Fig 4). The same analysis with late stage OA samples indicated that co-expression between the proteins in these pairs was largely lost at this stage of the disease (mean slope, 0.05). The same pattern could be detected when comparing protein co-expression in all 406 proteins, or when only selecting the differentially expressed proteins in one of the group comparisons but not the other, or even in proteins that did not show differential expression between any of the different groups (Supplemental Fig. S3, Fig. S4 and Fig.S5).

**Fig 5.**
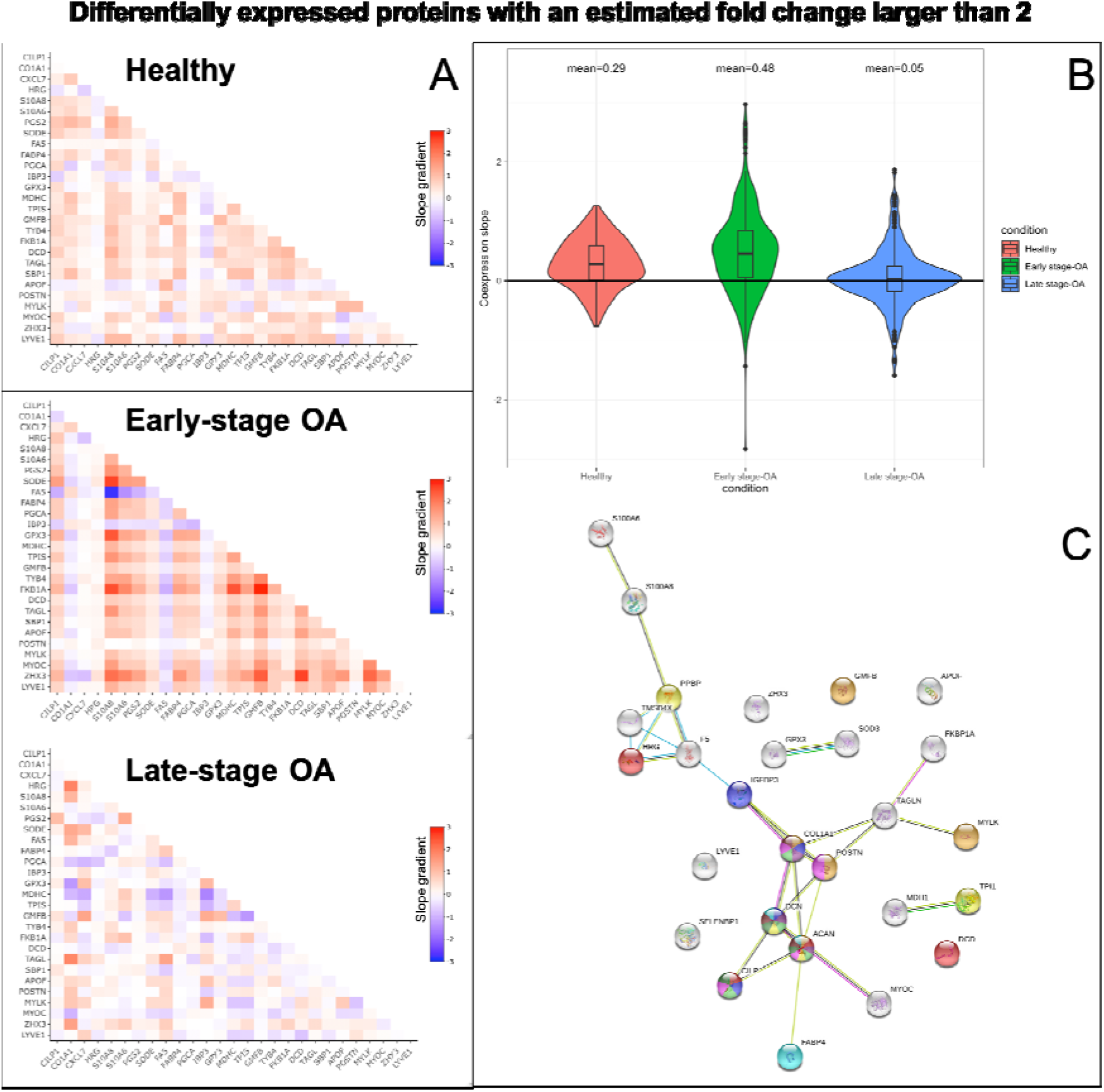
The estimated slope for each protein-protein co-expression pair between the 27 differentially expressed proteins with an estimated fold change larger than 2. A) The estimated slopes for each protein-protein co-expression pair of the 27 differentially expressed proteins with an estimated fold change larger than 2 are represented on the x and y axes for a for each group. B) The distribution of the estimated slopes for each of the protein–protein co-expression and the mean slope value is represented in the violin plot for each group. C) The protein-protein interaction network for the 27 proteins that were estimated to have a fold change larger than 2. Each color represents a previous study that suggested these proteins to be involved in osteoarthritis.

### Data quality

Repeatability coefficients estimated using the Bland-Altman approach for the 406 proteins included in the main analysis (i.e. the size of the difference between two measurements of the same sample) had (on a log_2_ scale) a median of 0.46, which corresponds to a fold change of 1.37. 75% of the proteins had a repeatability coefficient lower than 0.83 (corresponding to a fold change of 1.77).

### Validation of mass spectrometry label free quantification with Olink

Validating 36 differentially expressed proteins from the MS analysis with immuno-based proximity extension assay from Olink, a similar fold change pattern was found as in the label free MS results (Fig 5). Overall, the Olink data resulted in slightly higher fold changes than MS data. Olink data also had an overall lower dynamic range in comparison to the MS data.

**Fig 6.**
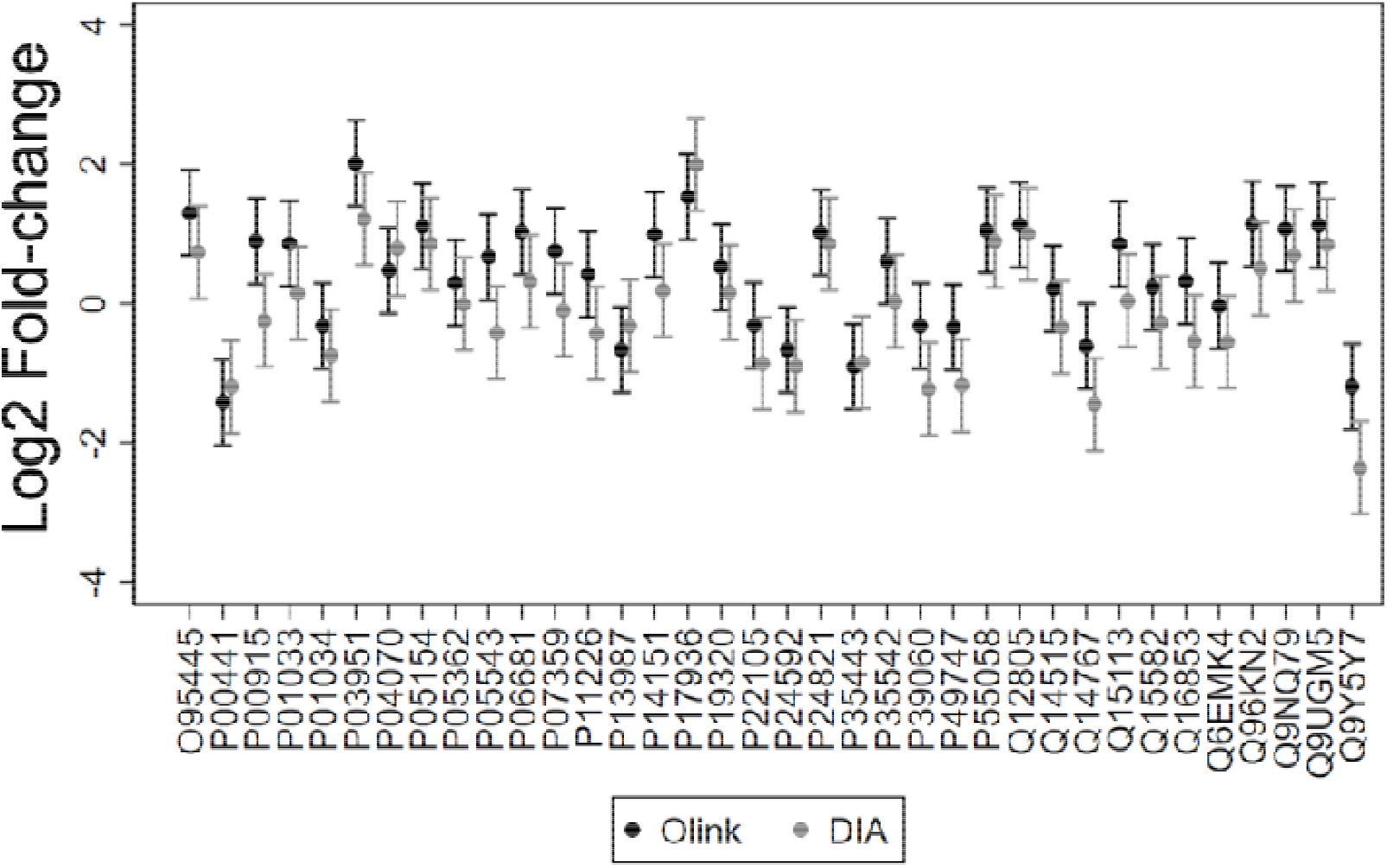
Validation of MS data. Comparison of fold changes for the late-stage OA vs controls between the two methods used, DIA and Olink, shown as differences with 95% CIs.

## Discussion

The results of this study generated evidence of a profound difference in the global protein profile of knee synovial fluid from controls compared to knee OA patients with more than 200 proteins differentially expressed between the different groups. Most of the proteins estimated to have a larger than 2-fold change between late-stage OA vs. controls and early-stage OA vs. controls were remarkably similar, suggesting their potential involvement in the OA disease process. The observed protein fold change patterns indicated an overall higher protein expression in early-stage OA, and a reduced global response in late-stage OA. Additionally, evaluation of the co-expression patterns of proteins within each of the three groups indicated that positive co-expression between proteins was increased in early-stage OA compared to controls, and that this co-expression was mostly lost in late-stage OA. The differential expression data revealed that similar pathways were detected to be activated or inhibited in the early stages OA and late stages OA in comparison to the controls. This indicates that the same types of mechanisms might be induced for degenerative meniscal tears as for late-stage OA.

The data were analysed both with an unsupervised dimensionality reduction test, PCA and through linear multilevel model^23–25^. A lot of the proteins that were differentially expressed in the statistical model were also among the principal components separating the different samples from each other, resulting in separated clusters in-between the different groups. Among the 27 proteins that were differentially expressed with a fold change larger than 2 have previously been reported to have a connection to osteoarthritis disease progression (Fig 4) ^13,26–31^. Proteins like cartilage intermediate layer protein 1 (CILP), aggrecan core protein (ACAN, PGCA), decorin (DCN, PGS2) and collagen type I (COL1A1) have been identified in multiple studies to be associated with OA. CILP, can act as an antagonist to TGFB1 mediated induction of cartilage matrix genes and IGF1-induced proliferation ^32^. Its overexpression may lead to impaired chondrocyte growth and matrix repair. The proteoglycan aggrecan (ACAN or PGCA) is a major component of the ECM of cartilaginous tissues ^33^. The major function of this protein is to resist compression in cartilage. It binds avidly to hyaluronic acid and forms negatively charged proteoglycan aggregates that attract water. Decorin (DCN) can affect the rate of collagen type I fibril formation, influencing the crosslinking of the ECM ^34^. These proteins exists in the molecular organization of normal articular cartilage and are present in the matrix surrounding chondrocytes^35^. In this study, all these three proteins were decreased in the synovial fluid of both early and late-stage OA in comparison to the controls. Periostin (POSTN) is associated with reparative processes involving cell adhesion and migration ^36^. It enhances incorporation of BMP1 in the fibronectin matrix of connective tissues, and subsequent proteolytic activation of lysyl oxidase for the crosslinking of the ECM. In this study, this protein was increased in both early and the late-stage OA in comparison to controls, and there was a 3-fold difference in its levels when comparing early and late-stage OA. Collagen type I (CO1A1) was found to be increased in the synovial fluid in early-stage OA but not in the late-stage. Additionally, higher expression of different types of collagens in early OA compared to the donor group, which might be an indicator of either ECM synthesis or degradation reflected in the synovial fluid. Among these, collagen type III (CO3A1) was increased in early OA but decreased in late-stage OA, compared to controls. Collagen III is known to increase during regeneration and wound healing ^37^, suggesting an attempted regeneration process in early OA. Taken together, the changes in these proteins suggest altered proliferation and development of chondrocytes, as well as widespread changes in the ECM especially in the early-stage of OA.

Some of the proteins that have a 2-fold or higher change in differential expression in OA patients compared to controls have previously not been reported to be associated with OA disease progression, e.g. ZHX3 and lymphatic vessel endothelial hyaluronic acid receptor 1 (LYVE1). ZHX3 acts as a transcriptional repressor involved in the early stages of mesenchymal stem cell osteogenic differentiation, which plays a fundamental role in skeletal tissue homeostasis ^38^. LYVE1 plays a role in autocrine regulation of cell growth and as a hyaluronan transporter, either mediating its uptake for catabolism within lymphatic endothelial cells themselves, or its transport into the lumen of afferent lymphatic vessels for subsequent re-uptake and degradation in lymph nodes ^39,40^. ZHX3 was increased in both early and late-stage OA while LYVE1 was decreased in the OA patients in comparison to the controls. The positive co-expression of these two proteins seen in controls and in early-stage OA, compared to the rest of the 27 differentially expressed proteins, seemed to be diminished in late-stage OA, suggesting that the crosstalk between these proteins is lost. This data could reflect a dysfunction in skeletal tissue homeostasis in late-stage OA.

Among the proteins that were found to be exclusively upregulated in late-stage OA was CRTAC1, a glycosylated ECM protein that can be found in the deep zone of articular cartilage ^41,42^. This protein is used as a marker to distinguish chondrocytes from osteoblasts and mesenchymal stem cells in culture. It can be detected in cartilage, bone, and cultured chondrocytes, but not in osteoblasts. The altered expression of this protein in the synovial fluid samples from late-stage OA cases might be a reflection of the severely damaged knee. It could also reflect higher exposure of the synovial fluid to the deep zone of articular cartilage. The transcription factors that were mapped through the differentially expressed proteins to be activated in the different OA stages were SRF and YBX1. SRF is involved in differentiation, proliferation and cell cycle progression ^43^. This suggests increased cellular activity in early-stage OA. YBX1 is involved in a number of cellular processes including proliferation, differentiation, and the cellular stress response ^44,45^. YBX1 activation in late-stage OA indicates the activation of responses for cell survival under stress conditions.

The degree of which inflammation contributes to OA pathogenesis and progression is an ongoing discussion in the field. There is still controversy whether it is predominantly an inflammatory, a low-grade inflammatory or a “wear and tear” mechanical driven disease ^46–48^. In this study, we have utilized discovery based MS which is capable of capturing a high dynamic range of molecules in a sample and we could not find a strong evidence of a typical inflammatory response associated with OA. On the other hand, this analysis may not detect the very low abundant inflammatory markers if no extensive fractionation is carried out. Well known inflammatory signalling molecules were only detected in low percentage, and the differential expression among disease stages of such molecules did not indicate that inflammation as the driving force in the OA pathogenesis. Previous studies supporting inflammation as a driving force of OA were not discovery based and only explored specific inflammatory molecules in bio fluids such as blood or plasma. ^46^ The lack of a robust inflammatory signature in our findings matches previous reports using discovery based proteomics ^13,15–17^. Furthermore, clinical trials that targeted inflammation such as IL antagonist therapies and anti-cytokine therapeutic drugs failed to reverse OA^49^. Although inflammatory molecules could be detected even at a low level, the degree of which they contribute to disease progression remains unknown.

Overall, these results show that a higher proportion of the detected proteins were upregulated in early-stage OA while late-stage OA was characterized by an imbalance of the homeostasis of the synovial fluid proteome. Furthermore, our data indicates that profiling the proteomic signature of a disease state may not be solely sufficient to elucidate the underlying mechanisms of OA. However, the disrupted cross talk between proteins across different stages of the disease might provide insights into the global mechanisms of disease onset and progression.

We would like to acknowledge some important limitations of this study. Four out of seven synovial fluid samples in the early-stage OA group were diluted during aspiration. To compensate for this, their total protein contents were normalized, leaving us with reduced individual variation detected in the differential expression analysis. On the other hand, the co-expression data were based on pairwise comparisons between proteins within the same sample, which is a major strength in this study. This minimizes any type of variation that could have come from the sample preparation or analysis. For the early-stage OA group, the same pattern of an overall increase in protein expression was detected in both the differential expression analysis and the co-expression analysis.

The samples used as controls was from post mortem donors. To our knowledge there is no study of how quickly death effects the proteome in synovial fluid. Forensic studies have investigated the change of specific biochemical like potassium, glucose and creatinine ^50^. Even though only very few metabolites showed a correlation between the concentration and time post-mortem the countable difference was only detected after 120 h. ^50,51^. In this study, all samples were obtained within 48 h post-mortem, and the specimens were frozen at −80°C within 2 h of extraction to be eligible as controls. Therefore, the proteomic status was assumed to be stable post-mortem.

## Conclusions

We found profound differences in the protein profiles of synovial fluid from controls and patients with different stages of OA. The synovial fluid of early-stage OA might represent a “raging battle field” of increased protein activity, while late-stage OA displays the “aftermath”. Further efforts should be made to elucidate the early processes in the disease, which have the greatest potential to be modified by therapeutic interventions.

## Acknowledgements

This work was supported by the European Research Council (ERC) under the European Union’s Horizon 2020 research and innovation programme (grant agreement #771121), the Swedish Research Council, the Foundation for Research in Rheumatology (FOREUM), the IAB Lundberg Foundation, the Greta and Johan Kock Foundation, the Swedish Rheumatism Association, the Österlund Foundation, the Governmental Funding of Clinical Research program within the National Health Service (ALF), the Royal Physiographic Society of Lund and the Faculty of Medicine, Lund University, Sweden. The funders had no role in study design, data collection and analysis, decision to publish, or preparation of the manuscript.

## Author contributions

Conception and design: NA, ME, and PÖ. Provision of study materials and tissue preparation: ME, NA, EF, VH, PÖ, JT, PN. Mass spectrometry analysis: NA. Statistical analysis: AT and NA. Interpretation of results: All co-authors. Drafting of the article: NA. Critical revision of the article for important intellectual content: NA, ME, AT, VH, PÖ and JT. Final approval of the article: All co-authors

