## Supplemental Figures for "Proteomics profiling of human synovial fluid suggests global increased protein interplay in early-osteoarthritis (OA) and lost in late-stage OA"

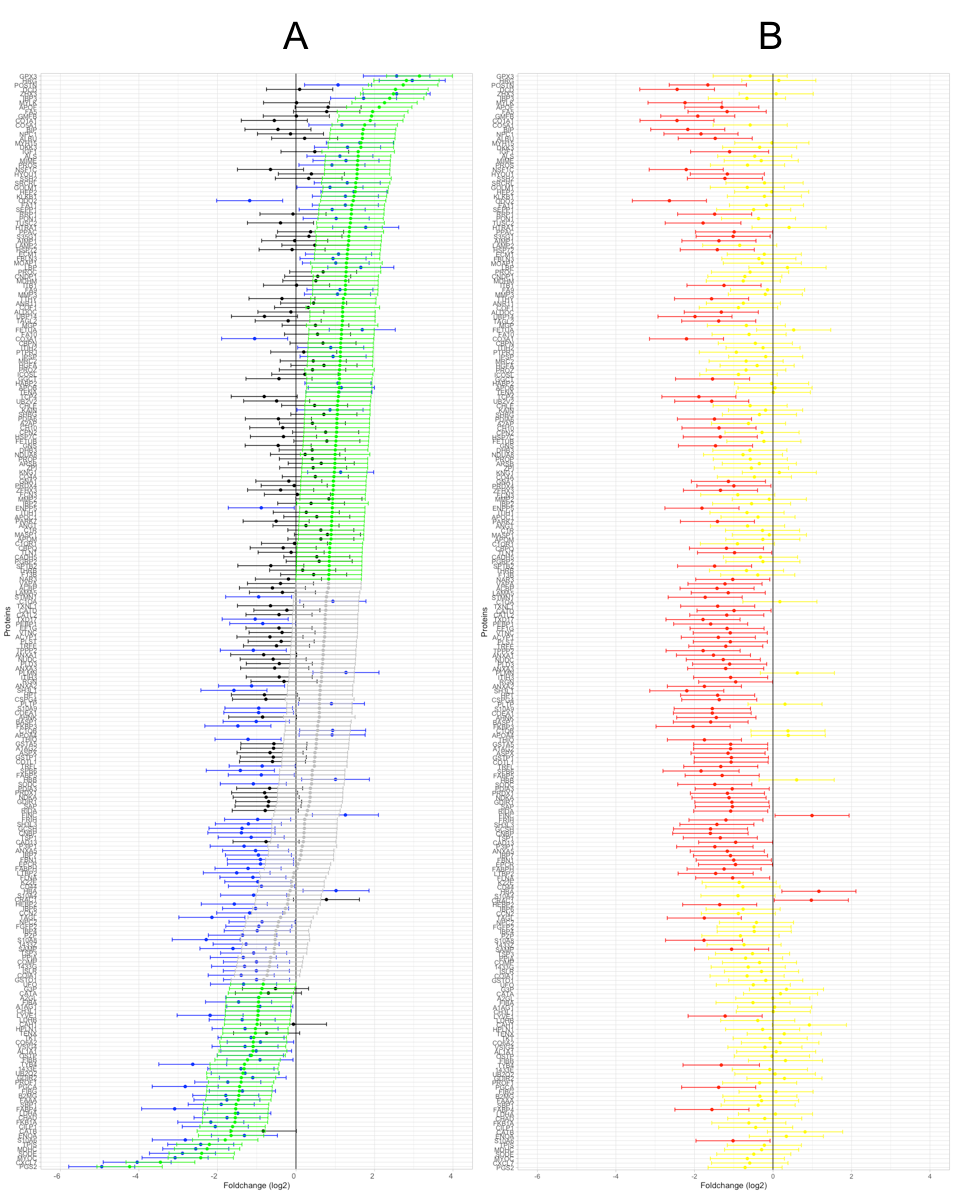

Fig. S1. Differentially expressed proteins with the estimated mean log_2_ fold change and the 95% CI. A) Comparison between early-stage OA vs controls (green and gray bars), Comparison between late-stage OA vs controls (blue and black bars). B) Comparison between late-stage OA vs early-stage OA (red and yellow bars). Blue, green, and red bars in their respective plots represent differentially expressed proteins whose 95% CIs exclude 0.

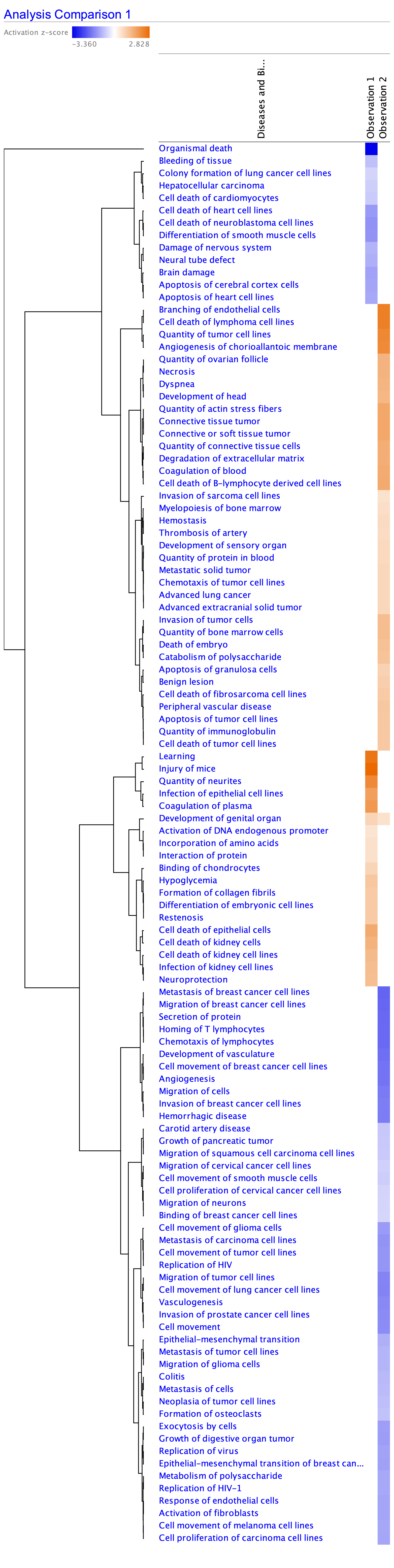

Fig. S2. The differentially expressed proteins are annotated and connected to different diseases and molecular functionalities coloured by the z-score (blue=down regulated and orange= up regulated). Observation 1: Diseases and molecular functionalities that are enriched in the early-stage OA group in comparison to controls. Observation 2: Diseases and molecular functionalities that are enriched in the late-stage OA group in comparison to donors.

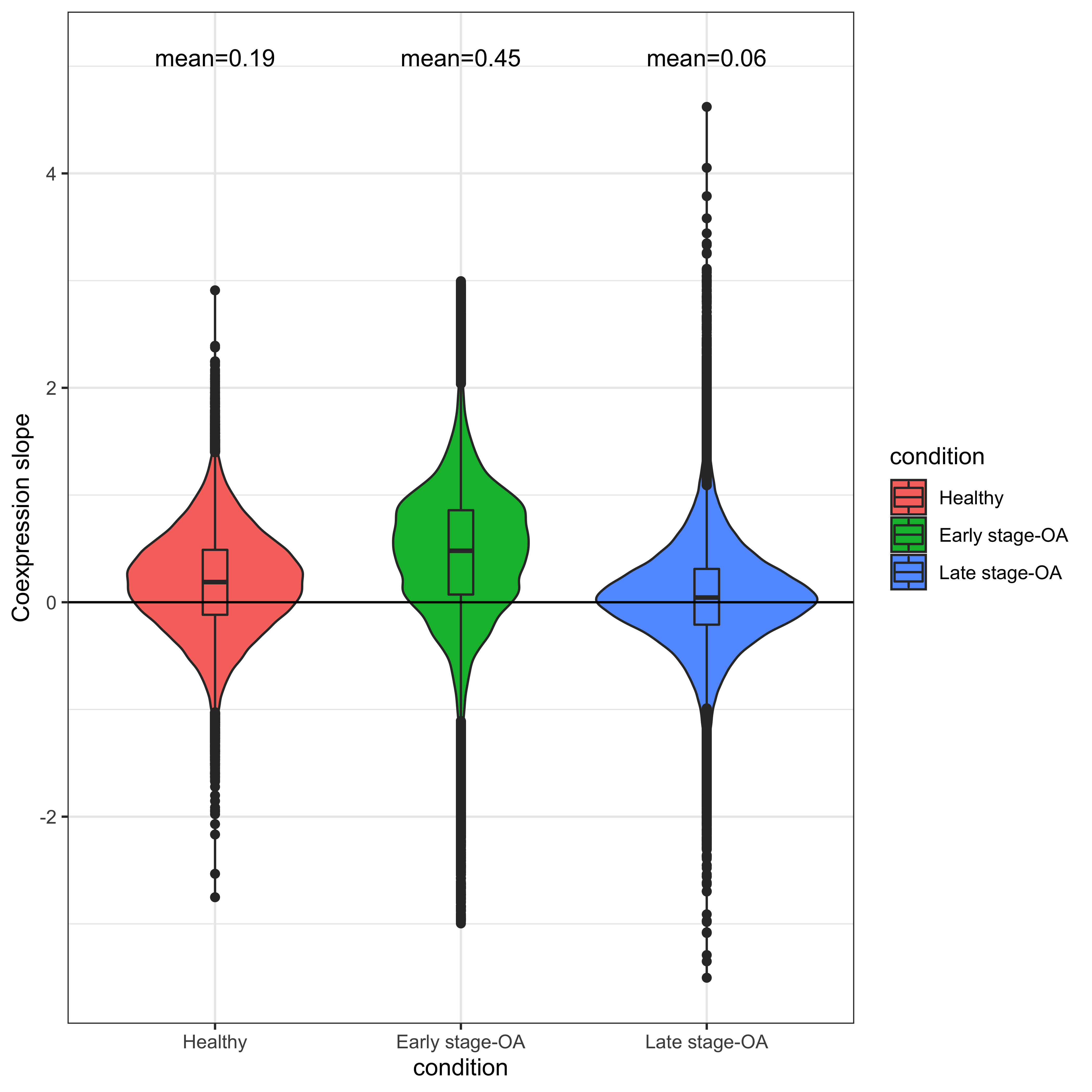

Fig. S3. Co-expression slope summery for 406 proteins. Represented in a violin plot with 95% CI boxplot and mean value for each group.

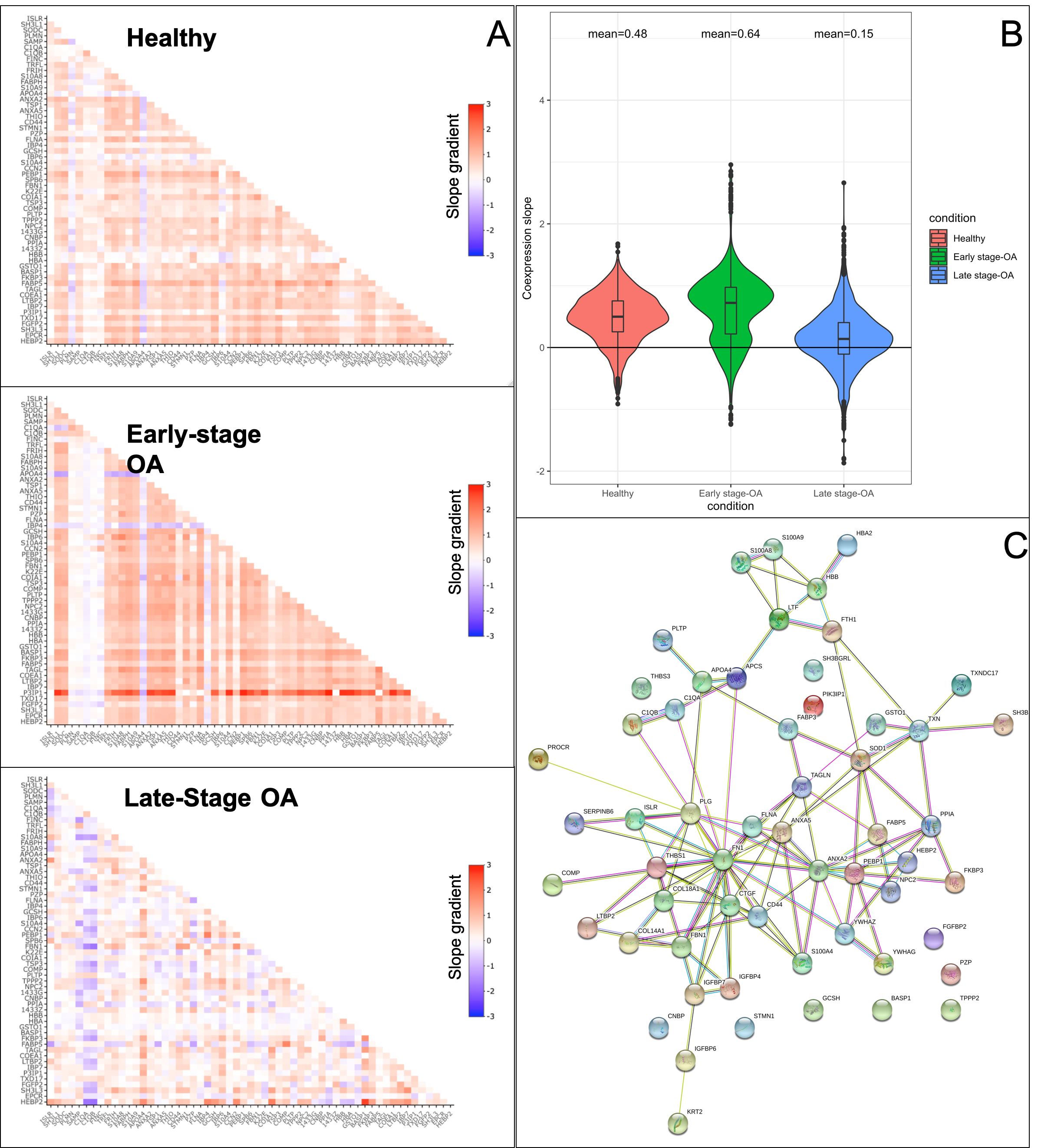

Fig. S4. **The estimated slope for each protein-protein co-expression pair between the differentially expressed proteins in the Late-stage OA vs Control comparison but were not differentially expressed in the Early-stage OA vs Control comparison.** A) The estimated slopes for each protein-protein co-expression pair are represented on the x and y axes for a for each group. B) The distribution of the estimated slopes for each of the protein–protein co-expression and the mean slope value is represented in the violin plot for each group. C) The protein-protein interaction network.

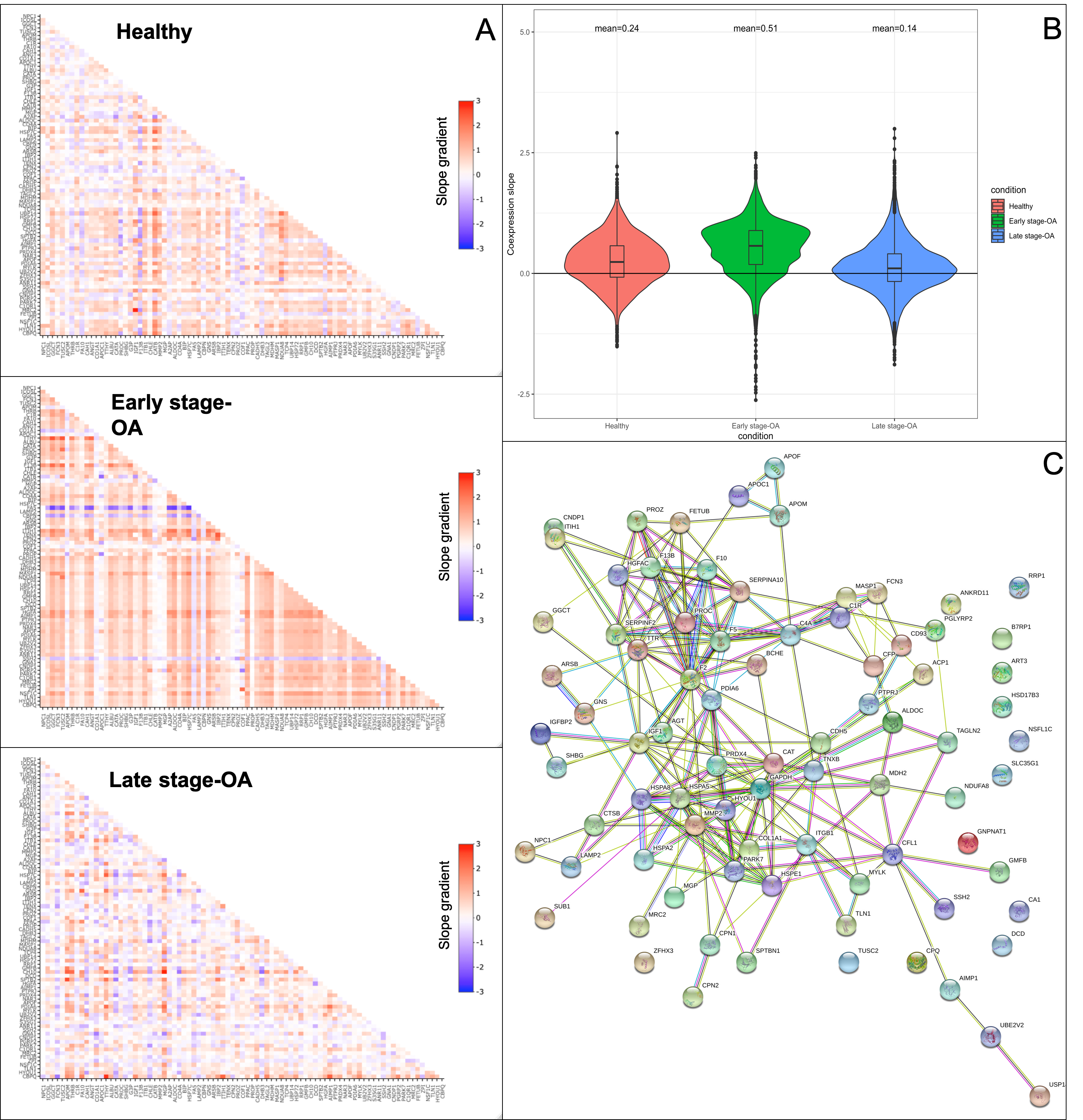

Fig. S5. **The estimated slope for each protein-protein co-expression pair between the differentially expressed proteins in the Early-stage OA vs Control comparison but were not differentially expressed in the Late-stage OA vs Control comparison.** A) The estimated slopes for each protein-protein co-expression pair are represented on the x and y axes for a for each group. B) The distribution of the estimated slopes for each of the protein–protein co-expression and the mean slope value is represented in the violin plot for each group. C) The protein-protein interaction network.
