## Supplementary material for "Proteomics profiling of human synovial fluid suggests global increased protein interplay in early-osteoarthritis (OA) and lost in late-stage OA": Supplemetal Table S1

**Table S1.** Estimated mean and 95% CI fold-change for 406 protein. Early-stage OA vs controls, Late-stage OA vs Contols and late-stage OA vs early-stage OA. Bold fold changes are differentially expressed protein with a 95% CI not crossing 1.

|  |  |  | Early-stage OA vs Controls | | Late-stage OA vs Controls | | Late-stage OA vs Early-stage OA | |
| --- | --- | --- | --- | --- | --- | --- | --- | --- |
| **Differentially expressed proteins** | Gene name | Protein AC | estimated mean | 95%CI | estimated mean | 95%CI | estimated mean | 95%CI |
|  | GPX3 | P22352 | **8,85** | **4,95-15,8** | **5,91** | **3,29-10,63** | 0,67 | 0,35-1,28 |
|  | HRG | P04196 | **7,03** | **3,95-12,51** | **7,79** | **4,35-13,95** | 1,11 | 0,58-2,13 |
|  | POSTN | Q15063 | **6,64** | **3,6-12,26** | **2,10** | **1,16-3,79** | **0,32** | **0,16-0,63** |
|  | DCD | P81605 | **5,79** | **3,25-10,32** | 1,07 | 0,59-1,91 | **0,18** | **0,1-0,36** |
|  | ZHX3 | Q9H4I2 | **5,59** | **3,13-9,99** | **5,91** | **3,27-10,68** | 1,06 | 0,55-2,04 |
|  | IBP3 | P17936 | **5,23** | **2,86-9,56** | **3,30** | **1,84-5,91** | 0,63 | 0,32-1,25 |
|  | MYLK | Q15746 | **4,77** | **2,68-8,5** | 1,01 | 0,56-1,81 | **0,21** | **0,11-0,41** |
|  | APOF | Q13790 | **4,35** | **2,44-7,74** | 1,76 | 0,98-3,16 | **0,41** | **0,21-0,78** |
|  | FA5 | P12259 | **3,87** | **2,07-7,22** | 1,72 | 0,96-3,09 | **0,44** | **0,22-0,89** |
|  | GMFB | P60983 | **3,79** | **2,13-6,75** | 1,01 | 0,56-1,8 | **0,27** | **0,14-0,51** |
|  | CO1A1 | P02452 | **3,71** | **2,09-6,61** | 0,68 | 0,38-1,22 | **0,18** | **0,1-0,35** |
|  | CO5A1 | P20908 | **3,36** | **1,89-5,98** | **2,25** | **1,26-4,03** | 0,67 | 0,35-1,29 |
|  | BIP | P11021 | **3,27** | **1,84-5,83** | 0,73 | 0,41-1,3 | **0,22** | **0,12-0,43** |
|  | NPC1 | O15118 | **3,24** | **1,82-5,76** | 0,91 | 0,51-1,63 | **0,28** | **0,15-0,54** |
|  | ALBU | P02768 | **3,23** | **1,81-5,75** | 1,16 | 0,65-2,09 | **0,36** | **0,19-0,69** |
|  | MYH15 | Q9Y2K3 | **3,15** | **1,76-5,63** | **3,09** | **1,71-5,58** | 0,98 | 0,51-1,89 |
|  | DKK3 | Q9UBP4 | **3,14** | **1,77-5,6** | **2,48** | **1,39-4,45** | 0,79 | 0,41-1,52 |
|  | IGF1 | P05019 | **3,00** | **1,58-5,69** | 1,39 | 0,77-2,53 | **0,46** | **0,23-0,93** |
|  | ALS | P35858 | **3,00** | **1,68-5,33** | **2,17** | **1,21-3,89** | 0,72 | 0,38-1,39 |
|  | MIME | P20774 | **2,97** | **1,67-5,29** | **2,42** | **1,35-4,33** | 0,81 | 0,42-1,56 |
|  | PROS | P07225 | **2,96** | **1,66-5,26** | **1,89** | **1,05-3,38** | 0,64 | 0,33-1,23 |
|  | NSF1C | Q9UNZ2 | **2,95** | **1,66-5,25** | 0,64 | 0,36-1,14 | **0,22** | **0,11-0,42** |
|  | HYOU1 | Q9Y4L1 | **2,94** | **1,65-5,23** | 1,31 | 0,73-2,35 | **0,45** | **0,23-0,86** |
|  | SSH2 | Q76I76 | **2,93** | **1,59-5,37** | 1,25 | 0,69-2,26 | **0,43** | **0,22-0,83** |
|  | SRCRL | A1L4H1 | **2,87** | **1,57-5,25** | **2,47** | **1,38-4,43** | 0,86 | 0,44-1,7 |
|  | GOLM1 | Q8NBJ4 | **2,87** | **1,61-5,11** | **1,82** | **1,02-3,26** | 0,64 | 0,33-1,22 |
|  | HEP2 | P05546 | **2,85** | **1,6-5,07** | **2,78** | **1,55-4,98** | 0,98 | 0,51-1,88 |
|  | KLKB1 | P03952 | **2,78** | **1,56-4,95** | **2,39** | **1,34-4,29** | 0,86 | 0,45-1,65 |
|  | ODO2 | P36957 | **2,75** | **1,54-4,89** | **0,44** | **0,25-0,79** | **0,16** | **0,08-0,31** |
|  | FA11 | P03951 | **2,68** | **1,51-4,77** | **2,39** | **1,33-4,28** | 0,89 | 0,46-1,71 |
|  | SEPP1 | P49908 | **2,66** | **1,5-4,74** | **1,89** | **1,06-3,39** | 0,71 | 0,37-1,37 |
|  | RRP1 | P56182 | **2,66** | **1,49-4,73** | 0,95 | 0,53-1,7 | **0,36** | **0,19-0,69** |
|  | PON1 | P27169 | **2,62** | **1,47-4,67** | **2,03** | **1,13-3,64** | 0,77 | 0,4-1,49 |
|  | TUSC2 | O75896 | **2,61** | **1,44-4,75** | 0,76 | 0,42-1,36 | **0,29** | **0,15-0,57** |
|  | HTRA1 | Q92743 | **2,58** | **1,45-4,59** | **3,42** | **1,91-6,13** | 1,33 | 0,69-2,55 |
|  | PPAC | P24666 | **2,57** | **1,4-4,7** | 1,29 | 0,72-2,32 | **0,50** | **0,25-1** |
|  | S35G1 | Q2M3R5 | **2,54** | **1,43-4,52** | 1,26 | 0,7-2,25 | **0,49** | **0,26-0,95** |
|  | AIMP1 | Q12904 | **2,54** | **1,43-4,52** | 0,98 | 0,55-1,75 | **0,39** | **0,2-0,74** |
|  | LAMP2 | P13473 | **2,51** | **1,41-4,47** | 1,40 | 0,78-2,5 | 0,56 | 0,29-1,07 |
|  | HSP72 | P54652 | **2,50** | **1,4-4,44** | 0,93 | 0,52-1,67 | **0,37** | **0,19-0,72** |
|  | ECM1 | Q16610 | **2,47** | **1,39-4,4** | **2,12** | **1,18-3,8** | 0,86 | 0,45-1,65 |
|  | FBLN3 | Q12805 | **2,47** | **1,39-4,39** | **1,92** | **1,07-3,44** | 0,78 | 0,41-1,5 |
|  | MOAP1 | Q96BY2 | **2,44** | **1,3-4,58** | **2,02** | **1,12-3,64** | 0,83 | 0,42-1,65 |
|  | LBP | P18428 | **2,43** | **1,33-4,45** | **3,14** | **1,75-5,63** | 1,29 | 0,65-2,55 |
|  | PROC | P04070 | **2,42** | **1,31-4,47** | 1,62 | 0,89-2,92 | 0,67 | 0,34-1,32 |
|  | CNDP1 | Q96KN2 | **2,41** | **1,35-4,29** | 1,47 | 0,82-2,63 | 0,61 | 0,32-1,17 |
|  | MDHM | P40926 | **2,40** | **1,35-4,27** | 1,42 | 0,79-2,54 | 0,59 | 0,31-1,14 |
|  | ITB1 | P05556 | **2,40** | **1,35-4,27** | 1,01 | 0,56-1,81 | **0,42** | **0,22-0,81** |
|  | FA9 | P00740 | **2,39** | **1,34-4,25** | **2,17** | **1,21-3,89** | 0,91 | 0,47-1,75 |
|  | MMP3 | P08254 | **2,38** | **1,34-4,24** | **2,08** | **1,16-3,73** | 0,87 | 0,45-1,68 |
|  | TTHY | P02766 | **2,31** | **1,3-4,11** | 0,78 | 0,44-1,4 | **0,34** | **0,18-0,65** |
|  | ANR11 | Q6UB99 | **2,30** | **1,29-4,1** | 1,36 | 0,76-2,45 | 0,59 | 0,31-1,14 |
|  | COF1 | P23528 | **2,28** | **1,19-4,37** | 1,24 | 0,69-2,25 | 0,54 | 0,27-1,09 |
|  | ALDOC | P09972 | **2,28** | **1,28-4,05** | 0,91 | 0,51-1,64 | **0,40** | **0,21-0,77** |
|  | UBP14 | P54578 | **2,27** | **1,28-4,04** | 0,57 | 0,32-1,03 | **0,25** | **0,13-0,49** |
|  | TAGL2 | P37802 | **2,27** | **1,28-4,04** | 0,87 | 0,49-1,56 | **0,38** | **0,2-0,74** |
|  | MGP | P08493 | **2,26** | **1,2-4,25** | 1,41 | 0,78-2,54 | 0,62 | 0,31-1,24 |
|  | FETUA | P02765 | **2,24** | **1,26-3,98** | **3,22** | **1,8-5,77** | 1,44 | 0,75-2,77 |
|  | FA10 | P00742 | **2,23** | **1,25-3,97** | 1,46 | 0,82-2,62 | 0,66 | 0,34-1,26 |
|  | CO3A1 | P02461 | **2,22** | **1,25-3,95** | **0,48** | **0,27-0,86** | **0,22** | **0,11-0,42** |
|  | CBPN | P15169 | **2,20** | **1,24-3,92** | 1,61 | 0,9-2,89 | 0,73 | 0,38-1,41 |
|  | ITIH2 | P19823 | **2,20** | **1,24-3,92** | **1,85** | **1,03-3,31** | 0,84 | 0,44-1,61 |
|  | PTPRJ | Q12913 | **2,19** | **1,23-3,9** | 1,15 | 0,64-2,05 | 0,52 | 0,27-1,01 |
|  | IPSP | P05154 | **2,19** | **1,23-3,89** | **1,92** | **1,07-3,45** | 0,88 | 0,46-1,69 |
|  | MRC2 | Q9UBG0 | **2,18** | **1,22-3,88** | 1,36 | 0,76-2,43 | 0,62 | 0,32-1,2 |
|  | HGFA | Q04756 | **2,17** | **1,22-3,86** | 1,64 | 0,91-2,94 | 0,76 | 0,39-1,45 |
|  | PROZ | P22891 | **2,15** | **1,17-3,98** | 1,34 | 0,74-2,41 | 0,62 | 0,31-1,25 |
|  | ICOSL | O75144 | **2,15** | **1,19-3,89** | 1,17 | 0,65-2,11 | 0,54 | 0,27-1,08 |
|  | GGCT | O75223 | **2,15** | **1,21-3,83** | 0,74 | 0,41-1,33 | **0,34** | **0,18-0,66** |
|  | HABP2 | Q14520 | **2,15** | **1,21-3,82** | **2,10** | **1,17-3,75** | 0,98 | 0,51-1,88 |
|  | APOB | P04114 | **2,15** | **1,21-3,82** | **2,23** | **1,24-3,99** | 1,04 | 0,54-1,99 |
|  | TENA | P24821 | **2,12** | **1,19-3,77** | **2,13** | **1,19-3,82** | 1,01 | 0,52-1,94 |
|  | TCP4 | P53999 | **2,11** | **1,19-3,76** | 0,57 | 0,32-1,02 | **0,27** | **0,14-0,52** |
|  | UB2V2 | Q15819 | **2,10** | **1,18-3,73** | 0,71 | 0,4-1,27 | **0,34** | **0,18-0,65** |
|  | CHLE | P06276 | **2,08** | **1,17-3,7** | 1,39 | 0,78-2,49 | 0,67 | 0,35-1,28 |
|  | KAIN | P29622 | **2,08** | **1,17-3,7** | **1,82** | **1,02-3,27** | 0,88 | 0,46-1,69 |
|  | SHBG | P04278 | **2,08** | **1,17-3,7** | 1,64 | 0,91-2,93 | 0,79 | 0,41-1,51 |
|  | PDIA6 | Q15084 | **2,06** | **1,16-3,67** | 0,73 | 0,41-1,31 | **0,36** | **0,19-0,68** |
|  | A2AP | P08697 | **2,06** | **1,16-3,67** | 1,34 | 0,75-2,39 | 0,65 | 0,34-1,25 |
|  | CH10 | P61604 | **2,05** | **1,15-3,65** | 0,79 | 0,44-1,42 | **0,39** | **0,2-0,74** |
|  | CPN2 | P22792 | **2,05** | **1,15-3,65** | 1,69 | 0,94-3,02 | 0,82 | 0,43-1,58 |
|  | HSP7C | P11142 | **2,03** | **1,14-3,62** | 0,80 | 0,45-1,44 | **0,39** | **0,21-0,76** |
|  | FETUB | Q9UGM5 | **2,03** | **1,14-3,6** | 1,72 | 0,96-3,09 | 0,85 | 0,44-1,64 |
|  | GNS | P15586 | **2,01** | **1,13-3,58** | 0,73 | 0,41-1,31 | **0,36** | **0,19-0,7** |
|  | DHB3 | P37058 | **2,00** | **1,12-3,56** | 1,33 | 0,74-2,38 | 0,67 | 0,35-1,28 |
|  | NDUA8 | P51970 | **1,99** | **1,08-3,68** | 1,18 | 0,64-2,17 | 0,59 | 0,29-1,18 |
|  | PROP | P27918 | **1,99** | **1,12-3,55** | 1,33 | 0,75-2,39 | 0,67 | 0,35-1,29 |
|  | ARSB | P15848 | **1,99** | **1,12-3,54** | 1,56 | 0,87-2,8 | 0,79 | 0,41-1,51 |
|  | ZPI | Q9UK55 | **1,98** | **1,11-3,54** | 1,35 | 0,75-2,44 | 0,68 | 0,35-1,31 |
|  | KNG1 | P01042 | **1,97** | **1,11-3,5** | **2,20** | **1,23-3,94** | 1,12 | 0,58-2,15 |
|  | CO4A | P0C0L4 | **1,95** | **1,1-3,47** | 1,41 | 0,79-2,52 | 0,72 | 0,37-1,39 |
|  | GNA1 | Q96EK6 | **1,94** | **1,09-3,45** | 0,88 | 0,49-1,58 | **0,46** | **0,24-0,88** |
|  | PRDX4 | Q13162 | **1,93** | **1,09-3,44** | 0,97 | 0,54-1,74 | **0,50** | **0,26-0,96** |
|  | ZFHX3 | Q15911 | **1,93** | **1,08-3,43** | 0,76 | 0,43-1,37 | **0,40** | **0,21-0,76** |
|  | FCN3 | O75636 | **1,91** | **1,07-3,4** | 1,03 | 0,57-1,84 | 0,54 | 0,28-1,03 |
|  | MMP2 | P08253 | **1,91** | **1,07-3,4** | 1,79 | 1-3,2 | 0,94 | 0,49-1,8 |
|  | IBP2 | P18065 | **1,90** | **1,01-3,58** | 1,31 | 0,72-2,35 | 0,69 | 0,34-1,36 |
|  | ENPP5 | Q9UJA9 | **1,90** | **1,07-3,38** | **0,54** | **0,3-0,97** | **0,29** | **0,15-0,55** |
|  | ITIH1 | P19827 | **1,90** | **1,07-3,38** | 1,20 | 0,67-2,15 | 0,63 | 0,33-1,22 |
|  | APOC1 | P02654 | **1,88** | **1,06-3,35** | 1,44 | 0,81-2,59 | 0,77 | 0,4-1,48 |
|  | PARK7 | Q99497 | **1,88** | **1,06-3,34** | 0,70 | 0,39-1,26 | **0,38** | **0,2-0,72** |
|  | ANGT | P01019 | **1,87** | **1,05-3,33** | 1,19 | 0,67-2,14 | 0,64 | 0,33-1,23 |
|  | C1R | P00736 | **1,87** | **1,05-3,32** | 1,55 | 0,87-2,78 | 0,83 | 0,43-1,6 |
|  | MASP1 | P48740 | **1,86** | **1,04-3,31** | 1,74 | 0,97-3,12 | 0,94 | 0,49-1,8 |
|  | APOM | O95445 | **1,86** | **1,04-3,3** | 1,55 | 0,87-2,78 | 0,84 | 0,43-1,61 |
|  | C1QR1 | Q9NPY3 | **1,82** | **1,03-3,25** | 0,97 | 0,54-1,74 | 0,53 | 0,28-1,03 |
|  | CBPQ | Q9Y646 | **1,81** | **1,02-3,23** | 0,80 | 0,45-1,43 | **0,44** | **0,23-0,85** |
|  | TLN1 | Q9Y490 | **1,81** | **1,02-3,22** | 0,92 | 0,51-1,64 | **0,51** | **0,26-0,98** |
|  | CADH5 | P33151 | **1,80** | **1,01-3,2** | 1,44 | 0,81-2,59 | 0,80 | 0,42-1,54 |
|  | PGRP2 | Q96PD5 | **1,80** | **1,01-3,2** | 1,51 | 0,84-2,7 | 0,84 | 0,44-1,61 |
|  | SPTB2 | Q01082 | **1,80** | **1,01-3,2** | 0,64 | 0,36-1,15 | **0,36** | **0,19-0,69** |
|  | THRB | P00734 | **1,79** | **1,01-3,19** | 1,12 | 0,63-2,02 | 0,63 | 0,33-1,21 |
|  | F13B | P05160 | **1,79** | **1-3,18** | 1,37 | 0,76-2,45 | 0,76 | 0,4-1,47 |
|  | NAR3 | Q13508 | **1,78** | **1-3,17** | 0,88 | 0,49-1,57 | **0,49** | **0,26-0,94** |
|  | VAPA | Q9P0L0 | 1,77 | 1-3,16 | 0,76 | 0,42-1,36 | **0,43** | **0,22-0,82** |
|  | ACBP | P07108 | 1,77 | 1-3,15 | 0,66 | 0,37-1,18 | **0,37** | **0,19-0,72** |
|  | LAMA5 | O15230 | 1,74 | 0,97-3,09 | 0,79 | 0,44-1,41 | **0,45** | **0,24-0,87** |
|  | STMN1 | P16949 | 1,71 | 0,96-3,05 | **0,52** | **0,29-0,93** | **0,30** | **0,16-0,58** |
|  | C1QA | P02745 | 1,70 | 0,95-3,02 | **1,92** | **1,07-3,44** | 1,13 | 0,59-2,17 |
|  | TXNL1 | O43396 | 1,69 | 0,95-3,01 | 0,64 | 0,36-1,14 | **0,38** | **0,2-0,72** |
|  | CATD | P07339 | 1,69 | 0,95-3,01 | 0,85 | 0,48-1,53 | **0,50** | **0,26-0,97** |
|  | CATL2 | O60911 | 1,68 | 0,94-2,98 | 0,74 | 0,41-1,33 | **0,44** | **0,23-0,85** |
|  | TXD17 | Q9BRA2 | 1,67 | 0,94-2,98 | **0,49** | **0,27-0,87** | **0,29** | **0,15-0,56** |
|  | PEBP1 | P30086 | 1,67 | 0,94-2,97 | **0,56** | **0,31-1** | **0,33** | **0,17-0,64** |
|  | EF1G | P26641 | 1,67 | 0,94-2,97 | 0,74 | 0,41-1,33 | **0,44** | **0,23-0,85** |
|  | VTNC | P04004 | 1,66 | 0,93-2,96 | 0,78 | 0,44-1,4 | **0,47** | **0,24-0,9** |
|  | ACYP1 | P07311 | 1,66 | 0,93-2,95 | 0,63 | 0,35-1,13 | **0,38** | **0,2-0,73** |
|  | PLST | P13797 | 1,63 | 0,92-2,91 | 0,77 | 0,43-1,38 | **0,47** | **0,24-0,91** |
|  | TRFE | P02787 | 1,63 | 0,92-2,9 | 0,71 | 0,4-1,27 | **0,43** | **0,23-0,84** |
|  | TPPP2 | P59282 | 1,62 | 0,91-2,88 | **0,47** | **0,26-0,84** | **0,29** | **0,15-0,56** |
|  | ANXA1 | P04083 | 1,62 | 0,91-2,88 | 0,57 | 0,32-1,01 | **0,35** | **0,18-0,67** |
|  | NUDC | Q9Y266 | 1,61 | 0,9-2,86 | 0,67 | 0,37-1,2 | **0,42** | **0,22-0,8** |
|  | PLD3 | Q8IV08 | 1,60 | 0,9-2,85 | 0,75 | 0,42-1,33 | **0,47** | **0,24-0,89** |
|  | ANXA3 | P12429 | 1,58 | 0,89-2,82 | 0,69 | 0,38-1,25 | **0,43** | **0,22-0,85** |
|  | PLMN | P00747 | 1,57 | 0,88-2,8 | **2,42** | **1,35-4,33** | 1,54 | 0,8-2,96 |
|  | ITIH3 | Q06033 | 1,57 | 0,88-2,8 | 0,74 | 0,42-1,33 | **0,47** | **0,25-0,91** |
|  | RGN | Q15493 | 1,56 | 0,88-2,78 | 0,81 | 0,45-1,45 | **0,52** | **0,27-1** |
|  | ANXA2 | P07355 | 1,53 | 0,86-2,73 | **0,46** | **0,25-0,82** | **0,30** | **0,15-0,57** |
|  | SH3L1 | O75368 | 1,53 | 0,86-2,73 | **0,33** | **0,19-0,6** | **0,22** | **0,11-0,42** |
|  | HPT | P00738 | 1,52 | 0,86-2,71 | 0,57 | 0,32-1,03 | **0,38** | **0,2-0,72** |
|  | CSPG4 | Q6UVK1 | 1,52 | 0,83-2,79 | 0,59 | 0,33-1,06 | **0,39** | **0,2-0,75** |
|  | PLTP | P55058 | 1,51 | 0,85-2,69 | **1,87** | **1,04-3,35** | 1,24 | 0,64-2,38 |
|  | S10A9 | P06702 | 1,51 | 0,85-2,69 | **0,52** | **0,29-0,94** | **0,34** | **0,17-0,67** |
|  | COEA1 | Q05707 | 1,51 | 0,85-2,69 | **0,52** | **0,28-0,96** | **0,34** | **0,17-0,68** |
|  | AHNK | Q09666 | 1,51 | 0,8-2,84 | 0,55 | 0,3-1,01 | **0,37** | **0,18-0,74** |
|  | BASP1 | P80723 | 1,49 | 0,84-2,65 | **0,50** | **0,28-0,89** | **0,33** | **0,17-0,64** |
|  | FKBP3 | Q00688 | 1,47 | 0,83-2,61 | **0,36** | **0,2-0,64** | **0,24** | **0,13-0,47** |
|  | C1QB | P02746 | 1,47 | 0,82-2,61 | **1,91** | **1,07-3,42** | 1,30 | 0,68-2,51 |
|  | APOA4 | P06727 | 1,45 | 0,81-2,57 | **1,89** | **1,06-3,39** | 1,31 | 0,68-2,52 |
|  | THIO | P10599 | 1,44 | 0,81-2,56 | **0,43** | **0,24-0,77** | **0,30** | **0,16-0,57** |
|  | GSTA5 | Q7RTV2 | 1,43 | 0,8-2,54 | 0,68 | 0,38-1,21 | **0,47** | **0,25-0,91** |
|  | A1AG2 | P19652 | 1,42 | 0,8-2,53 | 0,67 | 0,38-1,21 | **0,47** | **0,25-0,91** |
|  | ASPX | P26436 | 1,40 | 0,79-2,49 | 0,63 | 0,35-1,13 | **0,45** | **0,24-0,87** |
|  | GSTP1 | P09211 | 1,40 | 0,78-2,48 | 0,67 | 0,37-1,2 | **0,48** | **0,25-0,92** |
|  | COTL1 | Q14019 | 1,38 | 0,78-2,46 | 0,66 | 0,37-1,18 | **0,48** | **0,25-0,92** |
|  | TRFL | P02788 | 1,38 | 0,78-2,46 | **0,55** | **0,31-0,99** | **0,40** | **0,21-0,76** |
|  | SPB6 | P35237 | 1,33 | 0,75-2,37 | **0,37** | **0,21-0,68** | **0,28** | **0,14-0,55** |
|  | FABP5 | Q01469 | 1,33 | 0,75-2,37 | **0,54** | **0,3-0,97** | **0,41** | **0,21-0,78** |
|  | HBB | P68871 | 1,32 | 0,74-2,36 | **2,01** | **1,11-3,65** | 1,52 | 0,78-2,95 |
|  | SODC | P00441 | 1,32 | 0,74-2,34 | **0,47** | **0,26-0,84** | **0,36** | **0,19-0,69** |
|  | PDIA3 | P30101 | 1,29 | 0,72-2,3 | 0,63 | 0,35-1,12 | **0,49** | **0,25-0,94** |
|  | PRDX1 | Q06830 | 1,28 | 0,72-2,29 | 0,57 | 0,32-1,03 | **0,45** | **0,23-0,86** |
|  | NDKA | P15531 | 1,28 | 0,72-2,28 | 0,59 | 0,33-1,06 | **0,46** | **0,24-0,89** |
|  | GDIR1 | P52565 | 1,27 | 0,72-2,27 | 0,62 | 0,34-1,11 | **0,48** | **0,25-0,93** |
|  | SAP | P07602 | 1,25 | 0,7-2,23 | 0,61 | 0,34-1,09 | **0,49** | **0,25-0,94** |
|  | RIDA | P52758 | 1,23 | 0,69-2,18 | 0,58 | 0,32-1,04 | **0,47** | **0,25-0,91** |
|  | FINC | P02751 | 1,20 | 0,67-2,14 | **2,39** | **1,34-4,29** | **1,99** | **1,04-3,83** |
|  | FRIH | P02794 | 1,17 | 0,66-2,07 | **0,51** | **0,28-0,91** | **0,44** | **0,23-0,84** |
|  | SH3L3 | Q9H299 | 1,16 | 0,65-2,07 | **0,43** | **0,24-0,77** | **0,37** | **0,19-0,71** |
|  | GCSH | P23434 | 1,16 | 0,65-2,07 | **0,39** | **0,22-0,69** | **0,33** | **0,17-0,64** |
|  | CNBP | P62633 | 1,15 | 0,65-2,05 | **0,38** | **0,21-0,69** | **0,33** | **0,17-0,64** |
|  | TSP1 | P07996 | 1,15 | 0,65-2,05 | **0,45** | **0,25-0,81** | **0,39** | **0,21-0,76** |
|  | CAD13 | P55290 | 1,14 | 0,64-2,03 | 0,59 | 0,33-1,06 | **0,52** | **0,27-0,99** |
|  | P3IP1 | Q96FE7 | 1,12 | 0,61-2,05 | **0,40** | **0,22-0,73** | **0,36** | **0,18-0,7** |
|  | ANXA5 | P08758 | 1,10 | 0,62-1,95 | **0,49** | **0,27-0,88** | **0,45** | **0,23-0,86** |
|  | IBP7 | Q16270 | 1,09 | 0,61-1,94 | **0,52** | **0,29-0,92** | **0,47** | **0,25-0,91** |
|  | FBN1 | P35555 | 1,07 | 0,59-1,94 | **0,53** | **0,3-0,96** | **0,50** | **0,26-0,97** |
|  | EPCR | Q9UNN8 | 1,04 | 0,58-1,84 | **0,53** | **0,3-0,96** | **0,52** | **0,27-0,99** |
|  | FABPH | P05413 | 1,02 | 0,57-1,81 | **0,43** | **0,24-0,77** | **0,42** | **0,22-0,81** |
|  | LTBP2 | Q14767 | 0,97 | 0,53-1,77 | **0,35** | **0,19-0,63** | **0,36** | **0,19-0,7** |
|  | FLNA | P21333 | 0,95 | 0,54-1,7 | **0,47** | **0,26-0,84** | **0,49** | **0,26-0,94** |
|  | K22E | P35908 | 0,93 | 0,52-1,65 | **0,51** | **0,28-0,91** | 0,55 | 0,29-1,06 |
|  | CD44 | P16070 | 0,92 | 0,52-1,65 | **0,54** | **0,3-0,97** | 0,59 | 0,31-1,13 |
|  | HBA | P69905 | 0,90 | 0,51-1,6 | **2,03** | **1,13-3,63** | **2,25** | **1,17-4,32** |
|  | S10A4 | P26447 | 0,88 | 0,49-1,56 | **0,47** | **0,26-0,85** | 0,54 | 0,28-1,04 |
|  | CRAC1 | Q9NQ79 | 0,87 | 0,49-1,55 | 1,71 | 0,96-3,07 | **1,97** | **1,02-3,78** |
|  | HEBP2 | Q9Y5Z4 | 0,86 | 0,48-1,53 | **0,34** | **0,19-0,6** | **0,39** | **0,2-0,75** |
|  | IBP6 | P24592 | 0,83 | 0,47-1,48 | **0,49** | **0,27-0,88** | 0,59 | 0,31-1,13 |
|  | CCN2 | P29279 | 0,81 | 0,46-1,45 | **0,44** | **0,25-0,79** | 0,54 | 0,28-1,04 |
|  | TAGL | Q01995 | 0,76 | 0,43-1,36 | **0,23** | **0,13-0,41** | **0,30** | **0,15-0,57** |
|  | NPC2 | P61916 | 0,74 | 0,41-1,31 | **0,55** | **0,31-0,98** | 0,75 | 0,39-1,44 |
|  | FGFP2 | Q9BYJ0 | 0,72 | 0,41-1,29 | **0,52** | **0,29-0,93** | 0,72 | 0,37-1,38 |
|  | IBP4 | P22692 | 0,70 | 0,39-1,25 | **0,50** | **0,28-0,9** | 0,72 | 0,37-1,38 |
|  | PZP | P20742 | 0,69 | 0,38-1,27 | **0,39** | **0,21-0,71** | 0,56 | 0,28-1,11 |
|  | S10A8 | P05109 | 0,69 | 0,38-1,27 | **0,20** | **0,11-0,37** | **0,30** | **0,15-0,58** |
|  | 1433Z | P63104 | 0,69 | 0,39-1,23 | **0,41** | **0,23-0,74** | 0,60 | 0,31-1,15 |
|  | SAMP | P02743 | 0,68 | 0,38-1,21 | **0,33** | **0,18-0,59** | **0,48** | **0,25-0,93** |
|  | TSP3 | P49746 | 0,68 | 0,38-1,21 | **0,47** | **0,26-0,85** | 0,70 | 0,36-1,34 |
|  | PPIA | P62937 | 0,64 | 0,36-1,14 | **0,39** | **0,22-0,71** | 0,62 | 0,32-1,19 |
|  | COMP | P49747 | 0,63 | 0,36-1,12 | **0,50** | **0,28-0,89** | 0,79 | 0,41-1,52 |
|  | 1433G | P61981 | 0,62 | 0,35-1,11 | **0,40** | **0,23-0,72** | 0,65 | 0,34-1,24 |
|  | ISLR | O14498 | 0,61 | 0,34-1,08 | **0,50** | **0,28-0,89** | 0,82 | 0,42-1,57 |
|  | COIA1 | P39060 | 0,60 | 0,34-1,07 | **0,38** | **0,21-0,68** | 0,63 | 0,33-1,22 |
|  | GSTO1 | P78417 | 0,57 | 0,32-1,01 | **0,50** | **0,28-0,89** | 0,88 | 0,46-1,69 |
|  | UFO | P30530 | **0,56** | **0,32-1** | **0,40** | **0,22-0,71** | 0,71 | 0,37-1,36 |
|  | G3P | P04406 | **0,55** | **0,31-0,98** | 0,70 | 0,39-1,25 | 1,27 | 0,66-2,44 |
|  | CATA | P04040 | **0,54** | **0,3-0,96** | 0,61 | 0,34-1,1 | 1,14 | 0,59-2,2 |
|  | A2GL | P02750 | **0,52** | **0,29-0,92** | **0,52** | **0,29-0,93** | 1,00 | 0,52-1,92 |
|  | FIBA | P02671 | **0,52** | **0,29-0,92** | **0,36** | **0,2-0,65** | 0,70 | 0,37-1,35 |
|  | A1AG1 | P02763 | **0,51** | **0,29-0,91** | **0,53** | **0,29-0,95** | 1,03 | 0,54-1,98 |
|  | CH3L1 | P36222 | **0,51** | **0,29-0,91** | **0,51** | **0,29-0,92** | 1,01 | 0,52-1,93 |
|  | LYVE1 | Q9Y5Y7 | **0,51** | **0,29-0,91** | **0,22** | **0,12-0,39** | **0,43** | **0,22-0,83** |
|  | LDHB | P07195 | **0,51** | **0,28-0,9** | **0,39** | **0,22-0,69** | 0,76 | 0,4-1,47 |
|  | CAH1 | P00915 | **0,50** | **0,28-0,9** | 0,96 | 0,53-1,71 | 1,90 | 0,99-3,65 |
|  | HPLN1 | P10915 | **0,49** | **0,27-0,87** | **0,41** | **0,23-0,73** | 0,83 | 0,43-1,6 |
|  | TENX | P22105 | **0,49** | **0,27-0,87** | 0,60 | 0,33-1,07 | 1,22 | 0,63-2,34 |
|  | TKT | P29401 | **0,48** | **0,27-0,85** | **0,45** | **0,25-0,81** | 0,95 | 0,49-1,83 |
|  | CO6A2 | P12110 | **0,47** | **0,26-0,85** | **0,53** | **0,3-0,96** | 1,13 | 0,57-2,25 |
|  | VSIG4 | Q9Y279 | **0,47** | **0,26-0,83** | **0,41** | **0,23-0,73** | 0,87 | 0,45-1,67 |
|  | AL1A1 | P00352 | **0,47** | **0,26-0,85** | **0,50** | **0,27-0,92** | 1,06 | 0,53-2,13 |
|  | OSTP | P10451 | **0,46** | **0,25-0,83** | **0,45** | **0,25-0,8** | 0,98 | 0,51-1,91 |
|  | FIBB | P02675 | **0,43** | **0,24-0,76** | **0,53** | **0,3-0,95** | 1,25 | 0,65-2,4 |
|  | TYB4 | P62328 | **0,40** | **0,22-0,74** | **0,16** | **0,09-0,29** | **0,40** | **0,2-0,79** |
|  | 1433E | P62258 | **0,40** | **0,22-0,73** | **0,38** | **0,21-0,68** | 0,95 | 0,49-1,84 |
|  | UB2Q2 | Q8WVN8 | **0,39** | **0,21-0,74** | **0,41** | **0,22-0,74** | 1,04 | 0,51-2,11 |
|  | GDIR2 | P52566 | **0,38** | **0,21-0,7** | **0,47** | **0,26-0,84** | 1,22 | 0,63-2,37 |
|  | PROF1 | P07737 | **0,38** | **0,21-0,68** | **0,30** | **0,17-0,54** | 0,79 | 0,41-1,52 |
|  | PGCA | P16112 | **0,37** | **0,21-0,66** | **0,14** | **0,08-0,25** | **0,38** | **0,2-0,74** |
|  | FIBG | P02679 | **0,37** | **0,21-0,66** | **0,39** | **0,22-0,7** | 1,06 | 0,55-2,03 |
|  | B2MG | P61769 | **0,36** | **0,2-0,65** | **0,29** | **0,16-0,52** | 0,79 | 0,41-1,53 |
|  | FAAA | P16930 | **0,36** | **0,2-0,64** | **0,30** | **0,16-0,53** | 0,82 | 0,42-1,57 |
|  | SBP1 | Q13228 | **0,35** | **0,2-0,62** | **0,27** | **0,15-0,48** | 0,77 | 0,4-1,48 |
|  | FABP4 | P15090 | **0,34** | **0,19-0,61** | **0,12** | **0,07-0,21** | **0,34** | **0,18-0,65** |
|  | LDHA | P00338 | **0,34** | **0,19-0,61** | **0,36** | **0,2-0,64** | 1,04 | 0,54-2,01 |
|  | CHAD | O15335 | **0,34** | **0,19-0,61** | **0,30** | **0,17-0,53** | 0,86 | 0,45-1,66 |
|  | FKB1A | P62942 | **0,34** | **0,19-0,61** | **0,22** | **0,12-0,4** | 0,65 | 0,34-1,26 |
|  | CILP1 | O75339 | **0,33** | **0,18-0,58** | **0,24** | **0,13-0,43** | 0,74 | 0,38-1,42 |
|  | CATB | P07858 | **0,32** | **0,17-0,58** | 0,56 | 0,31-1,01 | 1,76 | 0,91-3,42 |
|  | ENOA | P06733 | **0,32** | **0,18-0,57** | **0,40** | **0,22-0,72** | 1,27 | 0,66-2,43 |
|  | S10A6 | P06703 | **0,29** | **0,16-0,51** | **0,14** | **0,08-0,25** | **0,49** | **0,26-0,95** |
|  | TPIS | P60174 | **0,22** | **0,12-0,39** | **0,19** | **0,1-0,33** | 0,86 | 0,45-1,65 |
|  | MDHC | P40925 | **0,21** | **0,12-0,37** | **0,17** | **0,1-0,31** | 0,82 | 0,43-1,57 |
|  | SODE | P08294 | **0,19** | **0,11-0,34** | **0,13** | **0,08-0,24** | 0,71 | 0,37-1,37 |
|  | MYOC | Q99972 | **0,19** | **0,1-0,33** | **0,12** | **0,07-0,21** | 0,64 | 0,33-1,22 |
|  | CXCL7 | P02775 | **0,09** | **0,05-0,17** | **0,06** | **0,03-0,11** | 0,66 | 0,34-1,3 |
|  | PGS2 | P07585 | **0,05** | **0,03-0,09** | **0,03** | **0,02-0,06** | 0,61 | 0,32-1,18 |
|  |  |  | Arthroscopy vs Donors | | Arthroplasty vs Donors | | Arthroplasty vs Arthroscopy | |
| **Not differentially expressed in any of the comparisons** | Gene name | Protein AC | estimated mean | 95%CI | estimated mean | 95%CI | estimated mean | 95%CI |
|  | MASP2 | O00187 | 1,19 | 0,67-2,11 | 1,01 | 0,57-1,81 | 0,85 | 0,44-1,64 |
|  | QSOX1 | O00391 | 1,43 | 0,8-2,55 | 1,13 | 0,63-2,02 | 0,79 | 0,41-1,52 |
|  | APOL1 | O14791 | 1,51 | 0,85-2,69 | 1,34 | 0,75-2,41 | 0,89 | 0,46-1,71 |
|  | TGON2 | O43493 | 1,15 | 0,64-2,04 | 0,68 | 0,38-1,22 | 0,59 | 0,31-1,14 |
|  | CD5L | O43866 | 0,72 | 0,4-1,32 | 0,80 | 0,45-1,44 | 1,11 | 0,57-2,15 |
|  | ATRN | O75882 | 1,48 | 0,83-2,63 | 1,10 | 0,61-1,96 | 0,74 | 0,39-1,42 |
|  | VNN1 | O95497 | 1,35 | 0,76-2,41 | 0,79 | 0,44-1,42 | 0,59 | 0,3-1,12 |
|  | GSHR | P00390 | 1,10 | 0,62-1,95 | 0,61 | 0,34-1,09 | 0,55 | 0,29-1,06 |
|  | CERU | P00450 | 0,86 | 0,48-1,54 | 0,85 | 0,47-1,52 | 0,99 | 0,51-1,9 |
|  | PGK1 | P00558 | 1,15 | 0,65-2,05 | 0,72 | 0,4-1,29 | 0,62 | 0,32-1,2 |
|  | CFAD | P00746 | 0,70 | 0,39-1,24 | 0,62 | 0,34-1,1 | 0,88 | 0,46-1,7 |
|  | FA12 | P00748 | 1,72 | 0,97-3,06 | 1,39 | 0,78-2,5 | 0,81 | 0,42-1,56 |
|  | CFAB | P00751 | 1,49 | 0,84-2,65 | 1,12 | 0,62-2 | 0,75 | 0,39-1,44 |
|  | ANT3 | P01008 | 1,13 | 0,64-2,02 | 1,02 | 0,57-1,83 | 0,90 | 0,47-1,73 |
|  | A1AT | P01009 | 1,04 | 0,58-1,85 | 0,70 | 0,39-1,25 | 0,67 | 0,35-1,29 |
|  | AACT | P01011 | 0,64 | 0,36-1,15 | 0,62 | 0,35-1,11 | 0,96 | 0,5-1,85 |
|  | A2MG | P01023 | 1,66 | 0,93-2,96 | 1,33 | 0,74-2,38 | 0,80 | 0,41-1,53 |
|  | CO3 | P01024 | 1,63 | 0,92-2,9 | 1,31 | 0,73-2,35 | 0,80 | 0,42-1,54 |
|  | CO5 | P01031 | 1,40 | 0,79-2,49 | 1,22 | 0,68-2,18 | 0,87 | 0,45-1,68 |
|  | TIMP1 | P01033 | 1,04 | 0,59-1,85 | 1,10 | 0,61-1,97 | 1,05 | 0,55-2,03 |
|  | CYTC | P01034 | 0,74 | 0,41-1,31 | 0,62 | 0,34-1,11 | 0,84 | 0,44-1,61 |
|  | IGF2 | P01344 | 1,66 | 0,93-2,96 | 1,57 | 0,88-2,82 | 0,95 | 0,49-1,82 |
|  | LMNA | P02545 | 1,08 | 0,61-1,93 | 0,60 | 0,34-1,08 | 0,55 | 0,29-1,07 |
|  | APOA1 | P02647 | 1,55 | 0,87-2,76 | 1,62 | 0,9-2,9 | 1,04 | 0,54-2 |
|  | APOE | P02649 | 1,51 | 0,85-2,7 | 1,36 | 0,76-2,43 | 0,90 | 0,47-1,72 |
|  | APOA2 | P02652 | 1,50 | 0,84-2,67 | 1,29 | 0,72-2,3 | 0,86 | 0,45-1,65 |
|  | APOC2 | P02655 | 1,07 | 0,6-1,91 | 1,09 | 0,61-1,96 | 1,02 | 0,53-1,96 |
|  | APOC3 | P02656 | 1,22 | 0,69-2,18 | 1,41 | 0,79-2,53 | 1,16 | 0,6-2,23 |
|  | C1QC | P02747 | 1,56 | 0,88-2,77 | 1,78 | 0,99-3,18 | 1,14 | 0,59-2,19 |
|  | CO9 | P02748 | 0,83 | 0,47-1,47 | 0,75 | 0,42-1,34 | 0,90 | 0,47-1,74 |
|  | APOH | P02749 | 1,25 | 0,7-2,23 | 1,13 | 0,63-2,02 | 0,90 | 0,47-1,73 |
|  | RET4 | P02753 | 1,38 | 0,78-2,46 | 1,32 | 0,74-2,37 | 0,96 | 0,5-1,84 |
|  | AMBP | P02760 | 0,95 | 0,53-1,69 | 0,96 | 0,54-1,72 | 1,01 | 0,53-1,95 |
|  | VTDB | P02774 | 1,05 | 0,59-1,86 | 1,03 | 0,58-1,85 | 0,99 | 0,51-1,9 |
|  | HEMO | P02790 | 1,04 | 0,59-1,86 | 0,95 | 0,53-1,7 | 0,91 | 0,47-1,75 |
|  | ALDOA | P04075 | 0,93 | 0,52-1,65 | 0,66 | 0,37-1,18 | 0,71 | 0,37-1,36 |
|  | LCAT | P04180 | 1,32 | 0,72-2,44 | 1,38 | 0,77-2,5 | 1,05 | 0,53-2,07 |
|  | A1BG | P04217 | 1,13 | 0,64-2,01 | 1,15 | 0,64-2,06 | 1,02 | 0,53-1,95 |
|  | K2C1 | P04264 | 1,16 | 0,65-2,07 | 0,62 | 0,35-1,12 | 0,54 | 0,28-1,03 |
|  | ALDOB | P05062 | 0,85 | 0,48-1,52 | 0,92 | 0,52-1,66 | 1,08 | 0,56-2,08 |
|  | APOD | P05090 | 0,89 | 0,5-1,59 | 0,89 | 0,5-1,59 | 1,00 | 0,52-1,92 |
|  | IC1 | P05155 | 0,98 | 0,55-1,75 | 0,97 | 0,54-1,74 | 0,99 | 0,51-1,9 |
|  | CFAI | P05156 | 1,43 | 0,8-2,54 | 1,20 | 0,67-2,15 | 0,84 | 0,44-1,62 |
|  | ICAM1 | P05362 | 1,05 | 0,59-1,86 | 1,08 | 0,6-1,94 | 1,04 | 0,54-1,99 |
|  | TETN | P05452 | 1,37 | 0,77-2,44 | 0,79 | 0,44-1,42 | 0,58 | 0,3-1,11 |
|  | THBG | P05543 | 0,91 | 0,51-1,62 | 0,79 | 0,44-1,41 | 0,87 | 0,45-1,67 |
|  | GELS | P06396 | 0,86 | 0,49-1,54 | 0,79 | 0,44-1,42 | 0,92 | 0,48-1,77 |
|  | CO2 | P06681 | 1,61 | 0,91-2,87 | 1,34 | 0,75-2,4 | 0,83 | 0,43-1,59 |
|  | CSF1R | P07333 | 0,96 | 0,51-1,82 | 1,02 | 0,56-1,85 | 1,06 | 0,53-2,11 |
|  | CO8A | P07357 | 1,43 | 0,81-2,55 | 1,31 | 0,73-2,35 | 0,92 | 0,48-1,76 |
|  | CO8B | P07358 | 1,20 | 0,67-2,13 | 1,07 | 0,6-1,93 | 0,90 | 0,47-1,72 |
|  | GP1BA | P07359 | 1,34 | 0,71-2,51 | 0,94 | 0,52-1,69 | 0,70 | 0,35-1,4 |
|  | CO8G | P07360 | 1,56 | 0,88-2,77 | 1,35 | 0,76-2,42 | 0,87 | 0,45-1,67 |
|  | TRY1 | P07477 | 1,50 | 0,84-2,67 | 0,89 | 0,49-1,59 | 0,59 | 0,31-1,14 |
|  | RNAS1 | P07998 | 1,07 | 0,6-1,91 | 0,69 | 0,39-1,24 | 0,65 | 0,34-1,24 |
|  | CO1A2 | P08123 | 1,27 | 0,7-2,33 | 0,88 | 0,49-1,57 | 0,69 | 0,35-1,36 |
|  | CBG | P08185 | 0,80 | 0,45-1,42 | 1,09 | 0,61-1,96 | 1,37 | 0,71-2,63 |
|  | CD14 | P08571 | 1,12 | 0,63-1,99 | 0,99 | 0,55-1,78 | 0,89 | 0,46-1,7 |
|  | CFAH | P08603 | 1,29 | 0,73-2,3 | 1,37 | 0,76-2,45 | 1,06 | 0,55-2,04 |
|  | VIME | P08670 | 1,16 | 0,65-2,06 | 1,22 | 0,68-2,19 | 1,05 | 0,55-2,03 |
|  | DOPO | P09172 | 1,30 | 0,73-2,32 | 0,92 | 0,49-1,71 | 0,70 | 0,35-1,4 |
|  | LEG1 | P09382 | 0,86 | 0,47-1,56 | 0,81 | 0,45-1,46 | 0,95 | 0,48-1,88 |
|  | SPRC | P09486 | 1,77 | 1-3,15 | 1,40 | 0,78-2,51 | 0,79 | 0,41-1,52 |
|  | GSTM1 | P09488 | 1,41 | 0,79-2,52 | 0,76 | 0,42-1,36 | 0,54 | 0,28-1,03 |
|  | C1S | P09871 | 1,63 | 0,92-2,91 | 1,75 | 0,98-3,13 | 1,07 | 0,56-2,06 |
|  | CO4B | P0C0L5 | 1,46 | 0,82-2,6 | 1,49 | 0,83-2,67 | 1,02 | 0,53-1,96 |
|  | CO7 | P10643 | 0,70 | 0,39-1,25 | 1,14 | 0,64-2,05 | 1,63 | 0,85-3,13 |
|  | CLUS | P10909 | 1,60 | 0,9-2,85 | 1,58 | 0,88-2,82 | 0,99 | 0,51-1,89 |
|  | MBL2 | P11226 | 1,26 | 0,71-2,24 | 0,77 | 0,43-1,38 | 0,61 | 0,32-1,17 |
|  | CO6A1 | P12109 | 0,89 | 0,5-1,58 | 1,30 | 0,72-2,32 | 1,46 | 0,76-2,81 |
|  | CO6A3 | P12111 | 1,55 | 0,87-2,75 | 0,92 | 0,51-1,64 | 0,59 | 0,31-1,14 |
|  | PEPD | P12955 | 0,79 | 0,44-1,4 | 0,67 | 0,38-1,21 | 0,85 | 0,44-1,64 |
|  | RINI | P13489 | 1,06 | 0,58-1,93 | 1,01 | 0,56-1,81 | 0,96 | 0,48-1,89 |
|  | CSPG2 | P13611 | 1,17 | 0,66-2,09 | 0,73 | 0,41-1,31 | 0,62 | 0,32-1,2 |
|  | K1C10 | P13645 | 0,98 | 0,55-1,75 | 0,57 | 0,32-1,02 | 0,58 | 0,3-1,11 |
|  | K2C5 | P13647 | 1,06 | 0,6-1,89 | 0,66 | 0,37-1,18 | 0,62 | 0,32-1,19 |
|  | CO6 | P13671 | 1,51 | 0,85-2,69 | 1,40 | 0,78-2,5 | 0,92 | 0,48-1,78 |
|  | PLSL | P13796 | 1,27 | 0,71-2,26 | 0,73 | 0,41-1,32 | 0,58 | 0,3-1,11 |
|  | CD59 | P13987 | 0,60 | 0,34-1,07 | 0,79 | 0,44-1,41 | 1,32 | 0,68-2,53 |
|  | LYAM1 | P14151 | 1,48 | 0,83-2,63 | 1,22 | 0,68-2,18 | 0,82 | 0,43-1,59 |
|  | FOLR2 | P14207 | 0,91 | 0,51-1,63 | 1,12 | 0,62-2 | 1,23 | 0,64-2,36 |
|  | KPYM | P14618 | 0,88 | 0,5-1,57 | 1,08 | 0,6-1,93 | 1,22 | 0,63-2,35 |
|  | AMPN | P15144 | 0,83 | 0,47-1,48 | 0,76 | 0,42-1,36 | 0,92 | 0,48-1,76 |
|  | TIMP2 | P16035 | 1,53 | 0,86-2,72 | 1,74 | 0,97-3,11 | 1,14 | 0,59-2,19 |
|  | AATC | P17174 | 1,55 | 0,87-2,76 | 1,06 | 0,58-1,91 | 0,68 | 0,35-1,32 |
|  | VINC | P18206 | 0,94 | 0,53-1,67 | 0,64 | 0,36-1,15 | 0,68 | 0,35-1,31 |
|  | PGAM1 | P18669 | 0,62 | 0,35-1,1 | 0,72 | 0,4-1,29 | 1,16 | 0,6-2,23 |
|  | VCAM1 | P19320 | 1,25 | 0,7-2,24 | 1,09 | 0,6-1,97 | 0,87 | 0,45-1,68 |
|  | GSTM3 | P21266 | 1,29 | 0,72-2,3 | 0,89 | 0,5-1,59 | 0,69 | 0,36-1,33 |
|  | PGS1 | P21810 | 1,06 | 0,6-1,89 | 0,83 | 0,46-1,49 | 0,78 | 0,41-1,51 |
|  | FBLN1 | P23142 | 0,81 | 0,46-1,45 | 0,57 | 0,32-1,03 | 0,70 | 0,37-1,35 |
|  | IBP5 | P24593 | 1,20 | 0,66-2,2 | 0,80 | 0,45-1,44 | 0,67 | 0,34-1,32 |
|  | ZA2G | P25311 | 1,00 | 0,56-1,78 | 0,80 | 0,45-1,43 | 0,80 | 0,42-1,54 |
|  | MOES | P26038 | 0,64 | 0,36-1,15 | 0,57 | 0,32-1,01 | 0,88 | 0,46-1,69 |
|  | HGFL | P26927 | 1,70 | 0,96-3,03 | 1,57 | 0,88-2,81 | 0,92 | 0,48-1,78 |
|  | IMPA1 | P29218 | 1,40 | 0,79-2,5 | 0,75 | 0,42-1,34 | 0,54 | 0,28-1,03 |
|  | PRDX6 | P30041 | 0,77 | 0,41-1,45 | 0,63 | 0,35-1,14 | 0,82 | 0,41-1,65 |
|  | PRDX2 | P32119 | 0,90 | 0,51-1,61 | 1,01 | 0,57-1,82 | 1,12 | 0,58-2,16 |
|  | MA1A1 | P33908 | 1,44 | 0,78-2,64 | 0,85 | 0,47-1,53 | 0,59 | 0,3-1,15 |
|  | RNAS4 | P34096 | 0,80 | 0,45-1,42 | 0,77 | 0,43-1,38 | 0,96 | 0,5-1,85 |
|  | TRY3 | P35030 | 1,60 | 0,9-2,85 | 0,83 | 0,46-1,49 | 0,52 | 0,27-1 |
|  | TSP4 | P35443 | 1,07 | 0,6-1,91 | 0,59 | 0,33-1,05 | 0,55 | 0,29-1,06 |
|  | K1C9 | P35527 | 1,18 | 0,67-2,11 | 0,80 | 0,45-1,44 | 0,68 | 0,35-1,3 |
|  | SAA4 | P35542 | 1,37 | 0,77-2,43 | 0,97 | 0,54-1,74 | 0,71 | 0,37-1,36 |
|  | PEDF | P36955 | 0,81 | 0,46-1,45 | 0,63 | 0,35-1,12 | 0,77 | 0,4-1,48 |
|  | FHR2 | P36980 | 0,70 | 0,39-1,24 | 1,01 | 0,56-1,8 | 1,44 | 0,75-2,77 |
|  | TALDO | P37837 | 1,26 | 0,71-2,25 | 0,78 | 0,44-1,4 | 0,62 | 0,32-1,19 |
|  | PTGDS | P41222 | 1,63 | 0,91-2,9 | 0,96 | 0,54-1,72 | 0,59 | 0,31-1,14 |
|  | MUC18 | P43121 | 1,44 | 0,81-2,56 | 0,90 | 0,5-1,62 | 0,63 | 0,33-1,21 |
|  | BTD | P43251 | 1,41 | 0,79-2,5 | 1,06 | 0,59-1,9 | 0,76 | 0,39-1,45 |
|  | AFAM | P43652 | 1,40 | 0,79-2,49 | 1,16 | 0,65-2,09 | 0,83 | 0,43-1,6 |
|  | GSTM5 | P46439 | 1,43 | 0,8-2,54 | 0,76 | 0,42-1,39 | 0,54 | 0,27-1,05 |
|  | TPC10 | P48553 | 0,94 | 0,53-1,67 | 0,78 | 0,44-1,4 | 0,83 | 0,43-1,6 |
|  | LUM | P51884 | 0,87 | 0,49-1,55 | 0,83 | 0,46-1,48 | 0,95 | 0,5-1,83 |
|  | CRIS3 | P54108 | 1,61 | 0,91-2,87 | 1,01 | 0,56-1,8 | 0,62 | 0,32-1,2 |
|  | AFAD | P55196 | 1,35 | 0,76-2,4 | 1,63 | 0,91-2,93 | 1,21 | 0,63-2,33 |
|  | LYSC | P61626 | 0,91 | 0,51-1,62 | 0,58 | 0,32-1,03 | 0,64 | 0,33-1,22 |
|  | PHLD | P80108 | 1,62 | 0,91-2,88 | 1,61 | 0,9-2,89 | 1,00 | 0,52-1,92 |
|  | PGBM | P98160 | 0,76 | 0,43-1,35 | 0,58 | 0,32-1,04 | 0,77 | 0,4-1,48 |
|  | FHR1 | Q03591 | 0,86 | 0,47-1,56 | 1,28 | 0,71-2,3 | 1,49 | 0,75-2,99 |
|  | KLC1 | Q07866 | 1,52 | 0,85-2,71 | 1,50 | 0,84-2,69 | 0,99 | 0,51-1,9 |
|  | LRP1 | Q07954 | 0,95 | 0,54-1,7 | 0,81 | 0,45-1,46 | 0,85 | 0,44-1,64 |
|  | FSTL1 | Q12841 | 1,57 | 0,88-2,79 | 1,18 | 0,66-2,12 | 0,75 | 0,39-1,45 |
|  | KGP2 | Q13237 | 1,63 | 0,92-2,9 | 1,15 | 0,64-2,07 | 0,71 | 0,37-1,36 |
|  | DAG1 | Q14118 | 1,02 | 0,56-1,87 | 1,06 | 0,59-1,91 | 1,04 | 0,53-2,05 |
|  | SPRL1 | Q14515 | 0,86 | 0,49-1,54 | 0,80 | 0,45-1,43 | 0,92 | 0,48-1,78 |
|  | ITIH4 | Q14624 | 1,58 | 0,89-2,81 | 1,21 | 0,68-2,17 | 0,77 | 0,4-1,47 |
|  | PCOC1 | Q15113 | 1,25 | 0,7-2,23 | 1,09 | 0,61-1,95 | 0,87 | 0,45-1,67 |
|  | BGH3 | Q15582 | 1,00 | 0,56-1,79 | 0,87 | 0,49-1,56 | 0,87 | 0,45-1,67 |
|  | ADIPO | Q15848 | 1,11 | 0,62-1,97 | 0,66 | 0,37-1,18 | 0,59 | 0,31-1,14 |
|  | DECR | Q16698 | 1,47 | 0,81-2,69 | 1,19 | 0,66-2,13 | 0,81 | 0,41-1,59 |
|  | AOC3 | Q16853 | 1,19 | 0,66-2,14 | 0,70 | 0,39-1,27 | 0,59 | 0,3-1,18 |
|  | VASN | Q6EMK4 | 1,02 | 0,57-1,82 | 0,70 | 0,39-1,25 | 0,68 | 0,36-1,31 |
|  | PXDC2 | Q6UX71 | 1,00 | 0,56-1,77 | 0,73 | 0,41-1,31 | 0,74 | 0,38-1,42 |
|  | PI16 | Q6UXB8 | 1,49 | 0,84-2,66 | 0,78 | 0,44-1,41 | 0,53 | 0,27-1,01 |
|  | CD109 | Q6YHK3 | 0,95 | 0,52-1,73 | 0,67 | 0,38-1,21 | 0,71 | 0,37-1,38 |
|  | KCNKI | Q7Z418 | 1,52 | 0,83-2,79 | 1,51 | 0,84-2,71 | 0,99 | 0,51-1,92 |
|  | K2C1B | Q7Z794 | 1,25 | 0,7-2,22 | 0,86 | 0,48-1,53 | 0,68 | 0,36-1,32 |
|  | TARSH | Q7Z7G0 | 0,93 | 0,52-1,65 | 0,62 | 0,35-1,11 | 0,67 | 0,35-1,29 |
|  | C163A | Q86VB7 | 0,56 | 0,32-1 | 0,67 | 0,38-1,21 | 1,20 | 0,62-2,3 |
|  | KLOTB | Q86Z14 | 1,15 | 0,65-2,05 | 0,95 | 0,53-1,71 | 0,83 | 0,43-1,6 |
|  | CILP2 | Q8IUL8 | 0,84 | 0,47-1,5 | 0,74 | 0,42-1,33 | 0,88 | 0,46-1,7 |
|  | DYH10 | Q8IVF4 | 0,96 | 0,51-1,79 | 1,10 | 0,59-2,05 | 1,15 | 0,58-2,27 |
|  | SCUB3 | Q8IX30 | 1,29 | 0,72-2,29 | 0,99 | 0,56-1,78 | 0,77 | 0,4-1,49 |
|  | PRG4 | Q92954 | 0,64 | 0,36-1,15 | 0,77 | 0,43-1,38 | 1,20 | 0,62-2,3 |
|  | CBPB2 | Q96IY4 | 1,58 | 0,89-2,82 | 1,42 | 0,79-2,54 | 0,90 | 0,47-1,72 |
|  | PEBP4 | Q96S96 | 0,84 | 0,47-1,5 | 0,66 | 0,37-1,18 | 0,78 | 0,41-1,5 |
|  | RARR2 | Q99969 | 1,12 | 0,63-1,99 | 1,53 | 0,85-2,74 | 1,36 | 0,71-2,62 |
|  | TBD2A | Q9BYX2 | 1,10 | 0,62-1,95 | 0,97 | 0,54-1,73 | 0,88 | 0,46-1,69 |
|  | MB12B | Q9H7P6 | 0,77 | 0,43-1,37 | 1,00 | 0,56-1,78 | 1,29 | 0,67-2,48 |
|  | CD248 | Q9HCU0 | 1,10 | 0,62-1,96 | 0,62 | 0,34-1,11 | 0,56 | 0,29-1,08 |
|  | IL1AP | Q9NPH3 | 1,57 | 0,88-2,79 | 0,89 | 0,5-1,59 | 0,57 | 0,3-1,09 |
|  | SPTN5 | Q9NRC6 | 1,51 | 0,82-2,76 | 0,89 | 0,5-1,6 | 0,59 | 0,3-1,17 |
|  | KRT82 | Q9NSB4 | 1,16 | 0,65-2,06 | 0,65 | 0,36-1,17 | 0,56 | 0,29-1,09 |
|  | C1RL | Q9NZP8 | 1,15 | 0,65-2,05 | 0,98 | 0,55-1,76 | 0,85 | 0,44-1,64 |
|  | CATZ | Q9UBR2 | 1,51 | 0,82-2,77 | 1,02 | 0,57-1,82 | 0,68 | 0,34-1,34 |
|  | BL1S6 | Q9UL45 | 0,85 | 0,46-1,57 | 0,67 | 0,37-1,2 | 0,78 | 0,4-1,55 |
|  | MINP1 | Q9UNW1 | 1,82 | 0,96-3,46 | 1,13 | 0,62-2,05 | 0,62 | 0,31-1,24 |
|  | NINL | Q9Y2I6 | 1,59 | 0,89-2,83 | 1,07 | 0,6-1,91 | 0,67 | 0,35-1,29 |
|  | FCGBP | Q9Y6R7 | 1,48 | 0,82-2,68 | 1,12 | 0,62-2,01 | 0,75 | 0,38-1,49 |
